# Glutamine Metabolism Supports α cell Mass and Glucagon Secretion

**DOI:** 10.64898/2026.07.09.735845

**Authors:** Anna Marie R. Schornack, Tyler J. Rodgers, Matthew Shou, Walter A. Siv, Linlin Yin, Katelyn Sellick, Varsha Chigurupati, Joshua Debo, Soham Saraf, Paula G. Nickles, Selina Park, Shannon E. Gibson, Nitin Shankar, Jordyn R. Dobson, Soma Behera, Jade E. Stanley, Angelina Ehara, Madushika Wimalarathne, Amber Crabtree, Austin Reuter, Alan Attie, Elma Zaganjor, Katie C. Coate, Yan Li, Jeffrey C. Rathmell, Mark P. Keller, David A. Jacobson, Wenbiao Chen, E. Danielle Dean

## Abstract

The liver-α cell axis is a finely tuned biological rheostat that regulates whole body amino acid availability. Pancreatic α cells secrete glucagon that regulates amino acid catabolism through gluconeogenesis and ureagenesis, yet the mechanisms linking amino acid levels to α cell growth and function are not fully understood. Here, we identify glutaminase, the enzyme that catalyzes glutamine catabolism, as a critical α cell regulator. Glutaminase is highly enriched in α cells across species. α cell expression of glutaminase is required for nutrient-dependent mTORC1 activation, suppression of AMPK signaling, and sustained expression of the glutamine transporter SLC38A5. This establishes a feed-forward loop linking glutamine metabolism to amino acid sensing and growth. Reduced glutaminase activity impairs dynamic glucagon secretion in response to low glucose and amino acids. Together, these findings highlight the importance of glutamine metabolism in α cell growth and hormone secretion and suggest it may play a role in α cell adaptation to hyperaminoacidemia.

## Introduction

Glucagon, a key regulator of nutrient metabolism, is secreted by pancreatic α cells, and plays a critical role in glucose homeostasis and intra-islet signaling for hormone regulation^1^. In a diseased state, the dysregulation of glucagon secretion can further exacerbate impaired glucose homeostasis^1^. Despite its critical role in maintaining metabolic homeostasis, the regulatory mechanisms governing glucagon secretion and α cell function remain underexplored, leaving significant gaps in our understanding of α cell dysregulation.

Recent studies have suggested that α cells may primarily function as amino acid sensors^2^. α cell activity is generally increased in postprandial metabolism, and the greatest response occurs when α cells are exposed to mixed nutrients^3^. Specifically, amino acids have been shown to be potent stimulators of glucagon secretion^2^. Circulating amino acid levels are strictly regulated by the liver, which controls their fractional uptake, synthesis, and catabolism^4^. Within the liver, glucagon receptor (GCGR) signaling leads to an upregulation of amino acid metabolism and hepatic glucose production^4^. While inhibiting the GCGR effectively reduces hyperglycemia, it also elicits hyperaminoacidemia, glucagon hypersecretion, and α cell hyperplasia in both humans and mice^4^. This phenomenon is attributed to a fundamental regulatory mechanism known as the liver-α cell axis^4^.

Amino acid tone is not static but is dynamically shaped across development and maturation as both hepatic metabolism and endocrine signaling capacity evolve. In early life, circulating amino acid levels are relatively elevated, reflecting high anabolic demand and an immature capacity for hepatic clearance^5^. As metabolic tissues mature, amino acid tone becomes more tightly constrained^5^, coinciding with enhanced efficiency of hepatic amino acid catabolism and increased fidelity of glucagon receptor signaling. Glucagon plays a critical role in this developmental refinement by linking amino acid availability to hepatic metabolic output, promoting their uptake and disposal while maintaining systemic balance^6^. Importantly, maturation is accompanied by improved sensitivity of α cells to amino acids and more robust downstream signaling through GCGR, allowing for tighter coupling between nutrient sensing and metabolic response^7^. These changes ensure that amino acid tone is appropriately matched to physiological demand, supporting growth early in life while preventing excess accumulation in adulthood.

L-glutamine has been identified as a critical amino acid in α cell proliferation and glucagon secretion^8^. While depletion of L-glutamine was shown to blunt amino acid-stimulated α cell proliferation^8^ *in vitro*, the exact mechanism remains unknown. In other biological contexts, most notably cancer, glutamine can become a conditionally essential nutrient that fuels a so-called “glutamine addiction” phenotype, in which cells rely disproportionately on glutamine to sustain growth and survival. This dependency arises because glutamine serves as both a carbon and nitrogen donor, supporting biosynthetic pathways, redox balance, and mitochondrial metabolism^9^. Following transport into the cell via amino acid transporters, glutamine is converted by glutaminase to glutamate, a key metabolic node that feeds into the tricarboxylic acid cycle and supports downstream signaling pathways such as mTORC1, a nutrient sensing hub that regulates cell growth^10^. The emergence of a similar glutamine-dependent program in α cells suggests that glutamine may function as a metabolic gatekeeper that enables proliferation in a manner that parallels well-established growth programs in other systems.

Despite the known role of glutamine metabolism in α cell proliferation and function, the precise molecular mechanisms linking glutamine metabolism to mTORC1 activation and glucagon secretion remain unclear. We propose that inhibiting glutaminase activity impairs α cell function by reducing mTORC1 activation and blunting glucagon secretion. Here, we identify glutaminase as a critical regulator of α cell nutrient sensing. We demonstrated that glutaminase-dependent glutamine metabolism is required for amino acid-stimulated glucagon secretion, mTORC1 activation, and adaptive α cell proliferation.

## Results

### Glutaminase is uniquely and robustly expressed in α cells across species

Previous studies show that high amino acid conditions induce α cell proliferation, with glutamine supporting this process by fueling metabolism and nucleotide biosynthesis, suggesting a key role for glutamine metabolism in α cell function^8^. To determine whether glutaminase, the enzyme that catalyzes the first step of glutamine catabolism, is enriched in α cells, we analyzed previously published single cell RNA sequencing data from human donors^11^. Kidney-type glutaminase (*GLS)* expression was highest in α cells, with levels ∼8.4-fold lower in β cells and ∼6.3-fold lower in δ cells (Figure 1A). This pattern was consistent with previously published mouse islet transcriptome data^12^, where α cells expressed *Gls* ∼3-fold higher than β and δ cells (Figure 1B), indicating a similar α cell enrichment of glutaminase in mice. Previous transcriptome data from zebrafish^13^ show that this α cell enrichment of glutaminase is consistent in zebrafish with a ∼4.1-fold higher expression of *glsb* in α cells compared to β cells and a ∼4.6-fold higher expression of *glsb* in α cells compared to δ cells (Figure 1C). This gene encodes the glutaminase enzyme that is found throughout the body, while the *glsa* gene is primarily found in the brain^14^. Liver-type glutaminase was lowly expressed in all pancreatic endocrine cells in humans (*GLS2),* mice (*Gls2*), and zebrafish (*gls2)* (Figure 1A-C).

**Figure 1.**
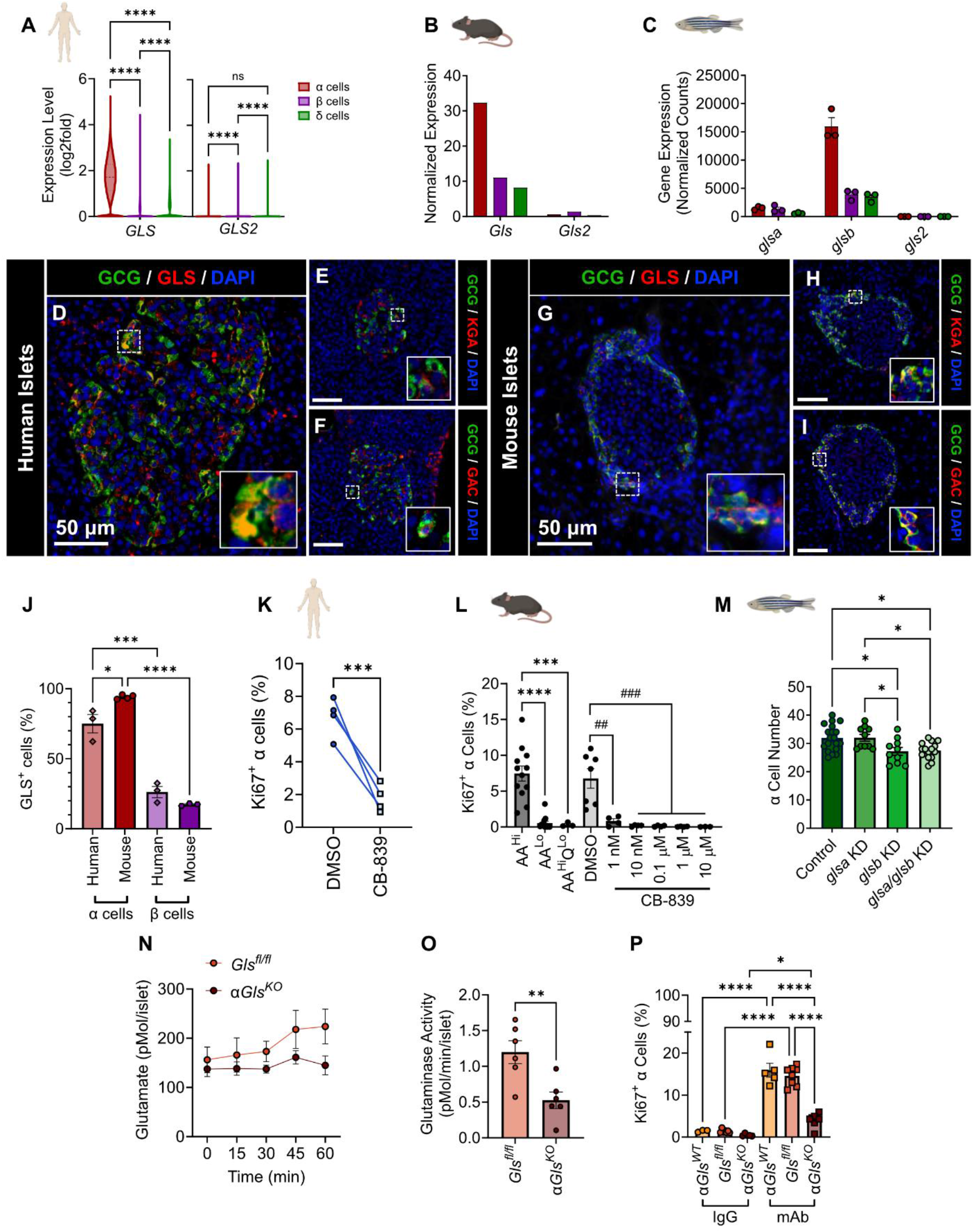
Glutaminase is an α cell identity marker that regulates amino acid-induced proliferation across species. (A) Comparison of *GLS* and *GLS2* expression in α (red), β (purple), and δ (green) cells from published human single cell RNA sequencing datasets, normalized to log2 fold change. One-way ANOVA, Tukey Post-Hoc. (B) Comparison of *Gls* and *Gls2* expression in α (red), β (purple), and δ (green) cells from published mouse transcriptome data set normalized to RPM. (C) Comparison of *glsa, glsb,* and *gls2* expression in α (red), β (purple), and δ (green) cells from published zebrafish transcriptome data set normalized to counts. (D-F) Representative immunostaining for glucagon and GLS1 (D), KGA (E), or GAC (F) in human islets (scale bar = 50 μm, inset width = 22 μm). (G-I) Representative immunostaining for glucagon and GLS1 (G), KGA (H), or GAC (I) in mouse islets (scale bar = 50 μm, inset width = 22 μm). (J) Quantification of GLS1 expression in α (red) and β (purple) in both humans and mice. (K) Quantification of α cell proliferation as determined by percent Ki67+/GCG+ cells per total GCG+ cells in human cadaveric donor islets treated with either DMSO or CB-839 (n= 4 each). (L) Ex vivo α cell proliferation analysis of isolated islets cultured in low amino acids, high amino acids, high amino acids without glutamine, or high amino acids with varying concentrations of CB-839. One-way ANOVA, Tukey Post-Hoc, Compared to AA^HI^, ***p<0.001, ****p<0.0001, Compared to DMSO, ^##^p<0.01, ^###^p<0.001. (M) Quantification of the number of α cells in zebra fish islets treated with either glsa, glsb, or glsa/glsb sgRNA or scramble control. One-Way ANOVA, Tukey Post-Hoc. (N) Islets from *Gls^fl/fl^*, and α*Gls^KO^* mice were lysed and incubated with 5 mM glutamine at 37°C. Reactions were heat-quenched at 0, 15, 30, 45, and 60 min, and glutamate production was measured. (O) Glutaminase activity was calculated from the slope of glutamate accumulation over time. Welch’s T-test. (P) Quantification of α cell proliferation as determined by percent Ki67+/GCG+ cells per total GCG+ cells in α*Gls^WT^, Gls^fl/fl^*, and α*Gls^KO^* mouse islets after two weeks treatment with GCGR mAb or control IgG (n=3-8 each).

Glutamine metabolism genes show clear cell-type-specific expression patterns within the pancreatic islet. α cells tend to express higher levels of genes involved in glutamine uptake and utilization (such as *Slc38* family transporters and *Gls*), consistent with a role for glutamine in supporting mTORC1 signaling, proliferation, and glucagon secretion (Extended Figure 1A-B). In contrast, β cells generally show lower reliance on glutamine catabolism (Extended Figure 1A-B) and instead favor glucose-centric metabolic pathways that couple glucose transport, glycolysis, and oxidative metabolism to insulin secretion^15^. δ cells exhibit an intermediate and more limited expression of glutamine metabolism genes (Extended Figure 1A-B), consistent with metabolic signaling that supports nutrient-stimulated somatostatin secretion and integration of paracrine inputs, rather than robust nutrient-driven anabolic or proliferative programs^16^.

Consistent with prior observations, we found that glutaminase expression was largely restricted to glucagon-positive α cells in both humans and mice (Figure 1D-J, Extended Figure 1C-D). Glutaminase is not as strongly expressed in β cells and δ cells (Extended Figure 1E-G). There are two isozymes of glutaminase, kidney-type (GLS) and liver-type (GLS2 or LGA), which are encoded by two separate genes^17^. Kidney-type glutaminase, which α cells primarily express^18^, consists of 19 exons and exists as multiple splice variants^17^. The first is kidney glutaminase (KGA), which contains exons 1-14 + 16-19, while Glutaminase C (GAC) includes exons 1-15. Human and mouse α cells express both glutaminase isoforms (Figure 1E-F, Figure 1H-I, Extended Figure 2A-D). In mice, α cells have higher exon usage of GAC than KGA (Extended Figure 1H)^19^.

To investigate the role of glutaminase in α cell metabolism, we first treated human islets with CB-839^20^, which is a selective pharmacological inhibitor for glutaminase. We found that inhibition of glutaminase blunts amino-acid induced α cell proliferation ∼4-fold (Figure 1K). In isolated mouse islets, CB-839 treatment using reduces α cell proliferation to a degree comparable to that observed under low amino acid conditions or when glutamine is selectively removed from high amino acid conditions (Figure 1L), highlighting the critical role of glutamine catabolism in α cell expansion. Upon genetic knockdown of *glsb* and *glsb/glsa* in zebrafish, we observe a decrease in the number of α cells (Figure 1M). These findings suggest that the role of glutaminase in regulating α cell proliferation may be conserved across vertebrate model organisms.

### Generation of pancreatic islet α cell-specific *Gls* knockout mice

α cells rely on glutamine both as a metabolic substrate and a signaling molecule^21^. This suggests that glutaminase may be a key regulator of α cell growth and nutrient-sensing pathways. To assess the importance of glutaminase in α cells, *Gls^fl/fl^* mice^22^ were crossed with Gcg-Cre^ERT2^ mice^23^ to generate offspring in which tamoxifen-induced Cre-mediated recombination excised exons 10 and 11 of *Gls* specifically in glucagon-expressing cells (Extended Figure 1I). Exons 10 and 11 encode the catalytic core common to both the KGA and GAC splice variants, so their excision effectively abolishes glutaminase enzymatic activity^24^. The tamoxifen-inducible Cre system also permits temporal control of gene deletion^25^, enabling us to examine glutaminase function in adult α cells independently of developmental effects. Littermate floxed mice lacking the Cre gene (*Gls^fl/fl^*) and Cre-positive mice lacking the floxed gene (α*Gls^WT^*) served as controls. Cre expression in α cells was validated via immunofluorescence (Extended Figure 2E-F). All mice were either treated with a GCGR monoclonal antibody (mAb) to interrupt glucagon signaling in the liver to elevated circulating amino acids or IgG antibody as a control.

Validation of the α*Gls^KO^*model demonstrated efficient and specific loss of *Gls1* expression in α cells compared to the floxed controls (*Gls^fl/fl^)* and the Cre controls (α*Gls^WT^*). Both the KGA and GAC isoforms were efficiently knocked out in the α cells of α*Gls^KO^* mice, and this loss was preserved under elevated amino acid conditions (Extended Figure 2C-F), demonstrating that the knockout is effective across splice variants and nutrient conditions. Indeed, only 5% of the α cells expressed GLS after tamoxifen treatment (Extended Figure 1C-D). The loss of *Gls* expression was maintained in α*Gls^KO^* mice treated with GCGR mAb, confirming the robustness of the knockout under conditions of amino acid abundance (Extended Figure 1D). Notably, the expression of glutaminase in δ cells significantly decreases under high amino acid levels, and this reduction is reversed with the loss of α cell-specific glutaminase (Extended Figure 1E and 1G).

### The loss of glutamine metabolism in α cells does not impact whole body physiology

We then sought to investigate the role of α cell-specific glutamine metabolism in whole body physiology. Loss of glutamine metabolism in α cells did not significantly affect body weight, blood glucose, or circulating amino acid responses to GCGR mAb treatment (Extended Figure 3A–D). These findings were consistent in both male and female mice, indicating that the observed phenotypes were not sex dependent.

Arginine stimulation increased circulating glucagon and insulin in all genotypes, although no significant genotype-dependent differences were observed (Extended Figure 3E–G). While *αGls^KO^* mice exhibited a greater fold increase in glucagon secretion, this trend did not reach statistical significance. Likewise, mixed-meal tolerance, glucose tolerance, and insulin tolerance tests revealed no significant differences between genotypes (Extended Figure 3H–M), indicating that α cell-specific glutaminase deficiency does not substantially alter whole-body glucose homeostasis under these conditions.

### Glutamine metabolism is required for α cell growth

To directly assess the functional consequence of α cell-specific *Gls* deletion of both isoforms, we measured glutaminase activity in isolated islets using a time-course assay. α*Gls^KO^* islets exhibited a marked reduction in glutamate production over time (Figure 1N), indicating a ∼2.28-fold decrease in glutaminase activity in α*Gls^KO^* islets relative to controls (Figure 1O). These results indicate that ∼56% of the glutaminase activity that occurs in islets comes from the α cells.

Elevated circulating amino acids are known to drive α cell proliferation^8^. We found that in control mice, amino acid-induced α cell proliferation, as quantified by Ki67-positive α cells, increased ∼12-fold under mAb treatment, consistent with a robust proliferative response to elevated amino acids (Figure 1P, Extended Figure 4A). In contrast, loss of glutaminase markedly reduced this response by ∼3.75-fold, demonstrating that glutamine metabolism is critical for amino acid-induced proliferation (Figure 1P, Extended Figure 4A). Both male and female mice were analyzed, and no sex-dependent differences were observed, so the data were pooled (Extended Figure 4B). Together, this data confirms that our α*Gls^KO^* model effectively reduces glutaminase activity in α cells and that glutamine metabolism plays a key role in regulating α cell proliferation.

### Glutamine metabolism controls α cell adaptative responses to amino acids

Elevated circulating amino acids are known to contribute to adaptive changes in islet architecture under conditions of disrupted glucagon signaling^8,26^. To assess how glutaminase impacts pancreatic islet cells, we first confirmed that all mice exhibited normal islet architecture and morphology regardless of amino acid levels (Figure 2A). Both male and female mice were included in all analyses, and no sex-dependent differences were observed in any measured parameter (Extended Figure 4C-H). Under elevated amino acid conditions, we found that the islet area is increased ∼1.5-fold in both control mice, but this effect is lost with the loss of α cell-specific glutaminase (Figure 2B). This appears to be because there is a larger number of smaller islets in the α*Gls^KO^* mouse model treated with GCGR mAb, while both control mice treated with GCGR mAb appear to have more mid-size islets. (Extended Figure 4H).

**Figure 2.**
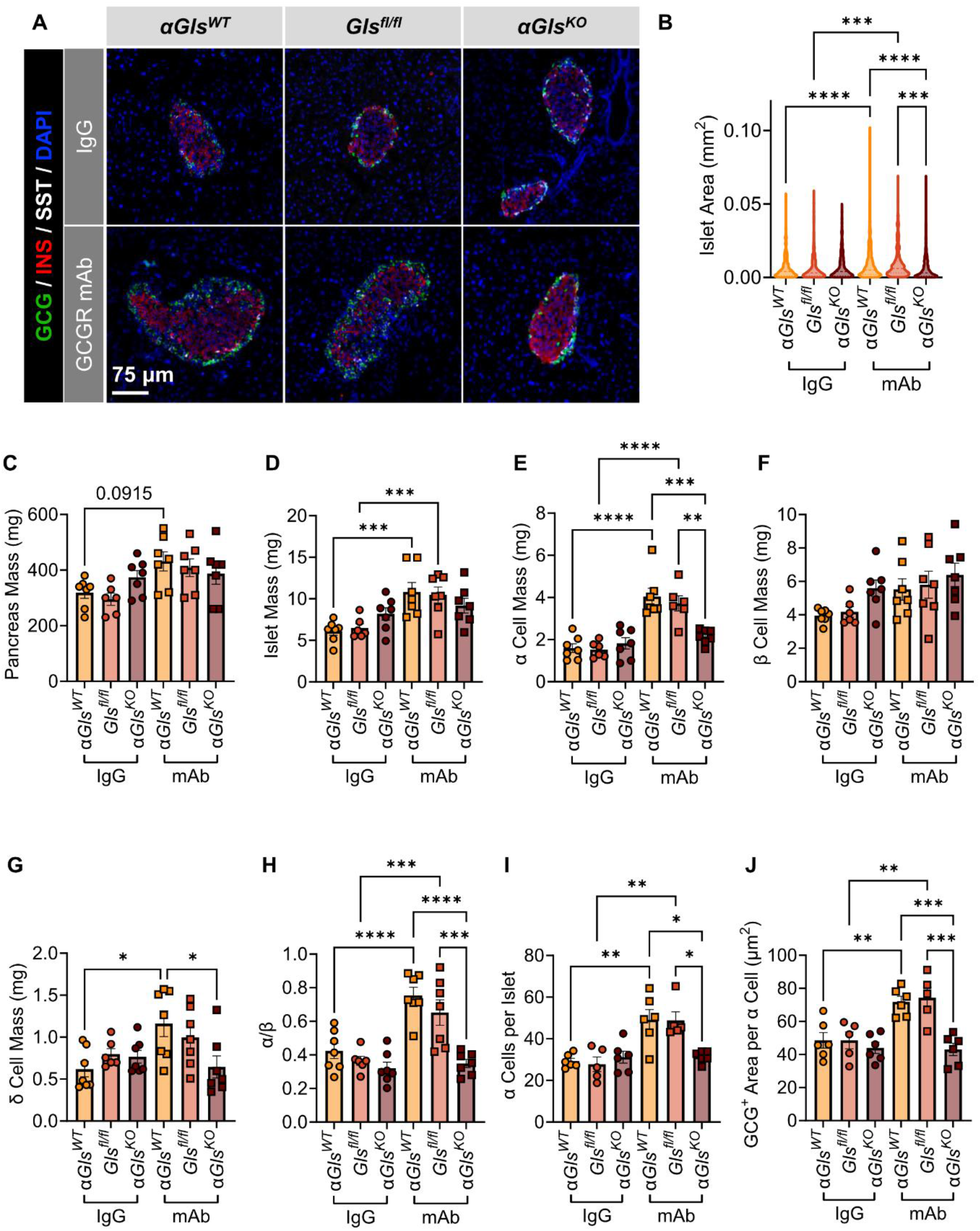
Glutamine metabolism supports α cell mass adaptations to circulating amino acids. (A) Representative immunostaining for glucagon, C-peptide, and somatostatin from *αGls^WT^, Gls^fl/fl^,* and α*Gls^KO^* mice treated with 10 days of IgG or mAb (scale bar = 75 μm). (B) Islet area as calculated from a mid-plane section. (C-G) Pancreas mass (C), islet mass (D), α cell mass (E), β cell mass (F), and δ cell mass (G) from α*Gls^WT^, Gls^fl/fl^*, and α*Gls^KO^* mice after two weeks of treatment with GCGR mAb or control IgG (n=6-7 each). (H) Calculated α to β cell ratio. (I) Quantification of the number of α cells per islet in a midplane section of the pancreas (n=5-6). (J) Quantification of the average size of an α cell in a midplane section of the pancreas (n=5-6).

To understand the changes in islet area, we assessed changes in cell mass. We first measured pancreas mass in α*Gls^WT^, Gls^fl/fl^,* and α*Gls^KO^* mice and found no significant differences across genotypes or amino acid conditions (Figure 2C). Under elevated amino acid conditions, total islet mass increased by ∼1.8-fold in α*Gls^WT^*and ∼1.6-fold in *Gls^fl/fl^* control mice, but not in the α*Gls^KO^* mice (Figure 2D). This expansion was primarily driven by a significant increase in α cell mass, which increased ∼2.3-fold in α*Gls^WT^*mice and ∼2.4-fold in *Gls^fl/fl^* mice (Figure 2E). The loss of glutaminase blunted the amino acid-induced α cell mass expansion by ∼2-fold, indicating that glutamine metabolism is required for a full α cell response to elevated amino acid levels (Figure 2E). Although not statistically significant, β cell mass showed a modest increase under normal amino acid conditions, suggesting that glutaminase-dependent glutamine metabolism has a more prominent role in α cell adaptation than in β cells (Figure 2F). δ cell mass is also elevated in response to high amino acids, but this effect is similarly reduced in α*Gls^KO^*mice, consistent with potential intercellular signaling or paracrine effects on δ cell expansion (Figure 2G). Unexpectedly, the α-to-β cell ratio increased under high amino acid conditions, and this shift was attenuated in the absence of glutaminase (Figure 2H).

Changes in α cell mass can result from altered proliferation or cell size^27^. Given that amino acids significantly altered α cell mass, we sought to determine whether this effect was due to hyperplasia or hypertrophy. We found that there was an increase in the number of α cells per islet by ∼1.7 fold in both control mice after mAb treatment. There was no increase in the number of α cells per islet in the α*Gls^KO^* mice (Figure 2I). There was also a ∼1.5-fold increase in the glucagon area per cell in both control mice after mAb treatment, which was blunted with the loss of α cell-specific glutaminase (Figure 2J). These findings demonstrate that glutaminase is required for amino acid–induced expansion of islet mass, through its effects on α cell hyperplasia and hypertrophy.

### Glutamine metabolism supports α cell mTORC1 signaling

To better understand the mechanisms underlying amino acid-induced α cell growth, we examined signaling pathways that regulate activation of mTORC1. We first evaluated if cell death was playing a role in the changes in α cell mass. We confirmed that apoptotic pathways were not activated by quantifying the expression of cleaved caspase 3 (Figure 3A-B). Given that amino acid–induced α cell proliferation, specifically through glutamine and arginine, is mTORC1-dependent^8,26^, we next assessed phosphorylated S6 protein (pS6), a downstream target of mTORC1, as a readout of nutrient-sensing signaling. We primarily measured phosphorylation at Ser240/244, a direct and specific target of S6K by mTORC1^28^. Consistent with increased proliferation, we observed that pS6 levels increased ∼10.4-fold in α*Gls^WT^*mice and ∼9.4-fold in *Gls^fl/fl^* mice following glucagon receptor blockade (Figure 3C, Extended Figure 5A). Loss of glutaminase blunted this activation by ∼2.35-fold, linking glutamine metabolism directly to nutrient-responsive mTORC1 signaling (Figure 3C, Extended Figure 5A). Both male and female mice were analyzed, and no sex-dependent differences were observed, therefore the data were pooled (Extended Figure 5B).

**Figure 3.**
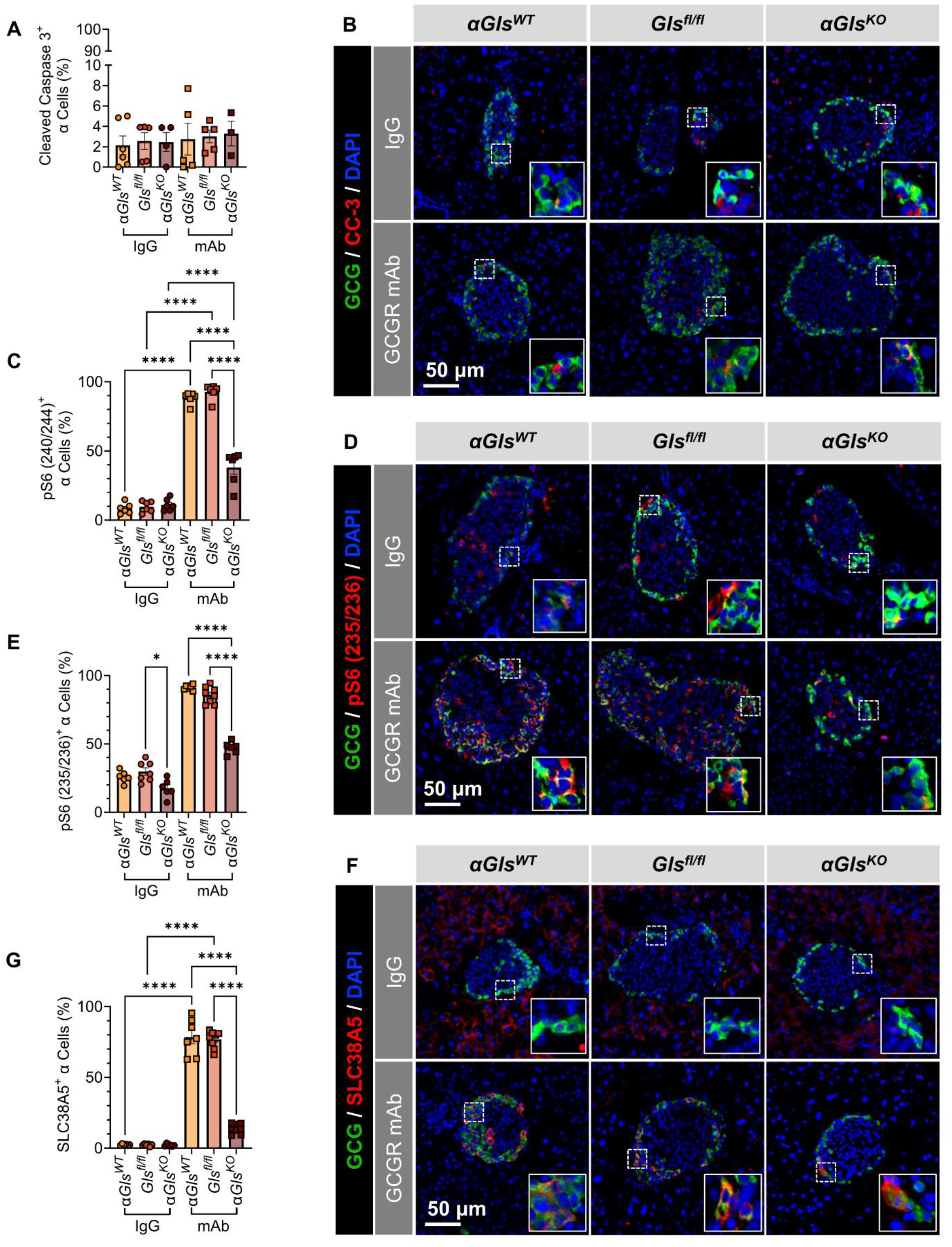
Glutamine metabolism supports α cell mTORC1 signaling. (A) Quantification of CC-3 expression in α cells as determined by CC-3^+^/GCG^+^ cells per total GCG^+^ cells in α*Gls^WT^, Gls^fl/fl^*, and α*Gls^KO^* mouse islets after treatment with GCGR mAb or control IgG (n=3-6). (B) Representative immunostaining for glucagon and cleaved caspase 3 (CC-3) from α*Gls^WT^, Gls^fl/fl^*, and α*Gls^KO^* mice (scale bar = 50 μm, inset width = 22 μm). (C) Quantification of mTORC1 activation in α cells as determined by percent pS6(240/244)+/GCG+ cells per total GCG+ cells in α*Gls^WT^, Gls^fl/fl^*, and α*Gls^KO^* mouse islets after two weeks treatment with GCGR mAb or control IgG (n=6 each). (D) Representative immunostaining for glucagon and pS6(235/236) from α*Gls^WT^, GGls^fl^lsfl^/fl^*, and α*Gls^KO^* mice (scale bar = 50 μm, inset width = 22 μm). (E) Quantification of S6 phosphorylation at Ser235/236 in α cells as determined by percent pS6(235/236)+/GCG+ cells per total GCG+ cells in α*Gls^WT^, Gls^fl/fl^*, and α*Gls^KO^* mouse islets after two weeks treatment with GCGR mAb or control IgG (n=5-6 each). (F) Representative immunostaining for glucagon and SLC38A5 from α*Gls^WT^, Gls^fl/fl^*, and α*Gls^KO^* mice (scale bar = 50 μm, inset width = 22 μm). (G) Quantification of SLC38A5 activation in α cells as determined by percent SLC38A5+/GCG+ cells per total GCG+ cells in α*Gls^WT^, Gls^fl/fl^*, and α*Gls^KO^* mouse islets after two weeks treatment with GCGR mAb or control IgG (n=6-8 each).

To evaluate whether amino acids activate broader growth signaling pathways beyond mTORC1 activation, we examined phosphorylation of S6 at Ser235/236. Phosphorylation at this site is a commonly used readout of nutrient-responsive signaling and reflects inputs beyond mTORC1 activity. We found that the loss of α cell-specific glutaminase resulted in a loss of phosphorylation of S6 at Ser235/236 under both physiological and elevated amino acid concentrations (Figure 3D-E). Phosphorylation at this residue suggests that α cells are integrating multiple growth signals^28^, indicating that amino acids may activate signaling pathways beyond those reflected by pS6 (Ser240/244).

Sustained growth signaling depends on continuous amino acid influx^29^, prompting us to examine whether glutaminase affects the expression of SLC38A5, a sodium-coupled neutral amino acid transporter that mediates glutamine uptake into α cells^30^. Consistent with previous findings^8,31,32^, SLC38A5 expression was strongly induced under elevated amino acid conditions (Figure 3F-G), paralleling increased mTORC1 activity. However, this induction was blunted ∼5.5-fold in α*Gls^KO^* mice (Figure 3F-G), indicating that a change in glutamine demand or utilization alters SLC38A5 expression in α cells. There was also no sex differences in the expression of SLC38A5 (Extended Figure 5C).

To assess how glutaminase influences amino acid–sensing pathways, we next examined the phosphorylation status of AMPK, AKT, and TSC2, key regulators of mTORC1 signaling under nutrient-limited conditions^29^. Under normal amino acid levels, α cells displayed high phosphorylated AMPK (pAMPK) expression, indicative of an energy- or nutrient-restricted state (Figure 4A-B). Elevated amino acid conditions markedly reduced pAMPK expression by ∼3.75-fold (Figure 4A-B), consistent with nutrient sufficiency and activation of mTORC1 signaling. Strikingly, this reduction was reversed by the loss of glutaminase, suggesting that glutamine metabolism contributes to the suppression of AMPK activity during amino acid abundance.

**Figure 4:**
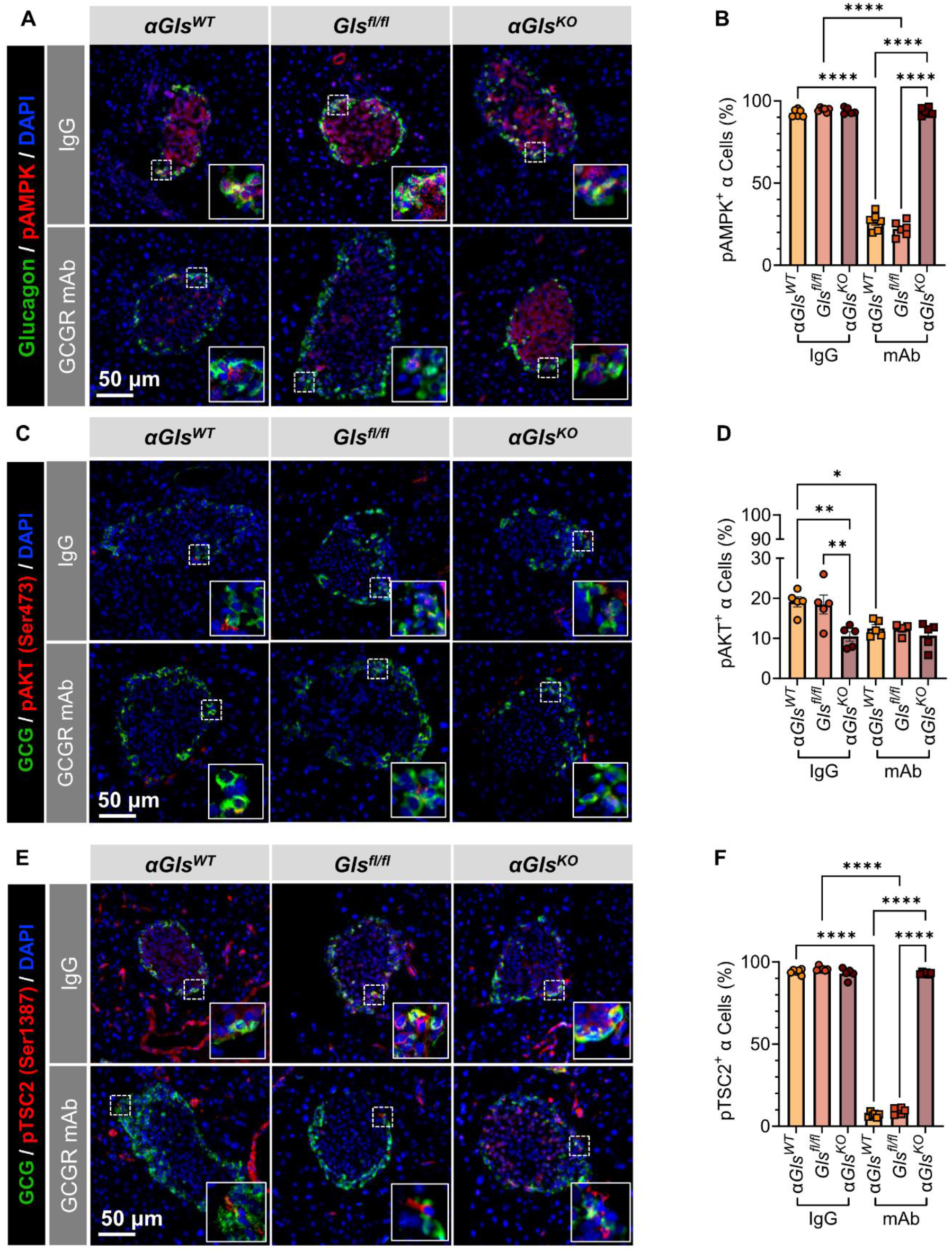
Glutamine metabolism supports AMPK-TSC2 inhibition in response to circulating amino acids. (A) Representative immunostaining for glucagon and pAMPK from α*Gls^WT^, Gls^fl/fl^*, and α*Gls^KO^* mice (scale bar = 50 μm, inset width = 22 μm). (B) Quantification of AMPK activation in α cells as determined by percent pAMPK+/GCG+ cells per total GCG+ cells in α*Gls^WT^, Gls^fl/fl^*, and α*Gls^KO^* mouse islets after two weeks treatment with GCGR mAb or control IgG (n=5-8 each). (C) Representative immunostaining for glucagon and pAKT from α*Gls^WT^, Gls^fl/fl^*, and α*Gls^KO^* mice (scale bar = 50 μm, inset width = 22 μm) (D) Quantification of AKT activation in α cells as determined by percent pAKT+/GCG+ cells per total GCG+ cells in α*Gls^WT^, Gls^fl/fl^*, and α*Gls^KO^* mouse islets after two weeks treatment with GCGR mAb or control IgG (n=4-5 each). (E) Representative immunostaining for glucagon and pTSC2 from α*Gls^WT^, Gls^fl/fl^*, and α*Gls^KO^* mice (scale bar = 50 μm, inset width = 22 μm). (F) Quantification of TSC2 activation in α cells as determined by percent pTSC2+/GCG+ cells per total GCG+ cells in α*Gls^WT^, Gls^fl/fl^*, and α*Gls^KO^* mouse islets after two weeks treatment with GCGR mAb or control IgG (n=5-6 each).

We next assessed phosphorylated AKT (pAKT) at Ser473, a direct target of mTORC2 that promotes mTORC1 activation through inhibition of TSC2^29^. Under normal amino acid conditions, pAKT levels were elevated (Figure 4C-D), reflecting active mTORC2 signaling in the nutrient-limited state. Elevated amino acid conditions reduced pAKT ∼1.5-fold in control mice (Figure 4D), likely due to mTORC1-dependent feedback inhibition. Notably, basal pAKT was blunted ∼1.8-fold in α*Gls^KO^* mice even under normal amino acid conditions, indicating that glutamine metabolism supports mTORC2 activity and AKT phosphorylation (Figure 4C-D).

TSC2 integrates inputs from both AMPK and AKT to regulate mTORC1^29^. We investigated phosphorylated TSC2 (pTSC2) at Ser1387, which is a direct target of AMPK, and observed similar regulatory pattern (Figure 4E-F). High amino acids reduced pTSC2 ∼10-fold in controls, whereas loss of glutaminase restored inhibitory TSC2 signaling. These findings are consistent with reactivation of the AMPK-TSC2 axis and attenuation of mTORC1 activation. Together, these findings reveal that glutaminase may integrate metabolic and signaling cues by suppressing AMPK–TSC2 activity while promoting amino acid transporter expression, thereby reinforcing nutrient-driven mTORC1 activation.

### Glutamine metabolism plays a crucial role in the relationship between amino acids and growth signaling in α cells

To determine whether the loss of glutaminase alters the relationship between amino acid availability and α cell growth signaling, we used linear regression models to quantify how these variables scale across conditions. As expected, amino acid levels were positively associated with α cell proliferation in both control mice, although the relationship explained a small fraction of the variance (R^2^ = 0.37 (α*Gls^WT^*), 0.3426 (*Gls^fl/fl^*)). Loss of α cell-specific glutamine metabolism attenuated the slope of this relationship, while increasing the proportion of variance explained by the model (R^2^ = 0.53) (Extended Figure 6A). There was a moderately strong positive relationship between amino acid levels and α cell mTORC1 activation in our control mice (R^2^ = 0.78 (α*Gls^WT^*), 0.60 (*Gls^fl/fl^*)). This relationship was weakened with the loss α cell-specific glutamine metabolism (R^2^ 0.40) (Extended Figure 6B). There was a positive relationship between α cell proliferation and α cell mTORC1 activation in all mice, but the proportion of variance was exceedingly small (R^2^ = 0.02 (α*Gls^WT^*), 0.02 (*Gls^fl/fl^*), 0.02 (α*Gls^KO^*)) (Extended Figure 6C). We found that both α cell proliferation (R^2^ = 0.77 (α*Gls^WT^*), 0.60 (*Gls^fl/fl^*)) and α cell mTORC1 activation (R^2^ = 0.67 (α*Gls^WT^*), 0.77 (*Gls^fl/fl^*)) had a strong positive relationship with α cell mass in our control mice, but the loss of α cell-specific glutamine metabolism attenuated both relationships (Ki67: R^2^ = 0.02; pS6 (240/244): R^2^ = 0.22) (Extended Figure 6D-E). Consistent with previous findings^32^, we see that there is a strong positive relationship between amino acid levels and SLC38A5 expression on α cells (R^2^ = 0.76 (α*Gls^WT^*), 0.76 (*Gls^fl/fl^*)). The loss of glutamine metabolism attenuated this relationship (R^2^ = 0.64) (Extended Figure 6F). These data implicate glutamine metabolism in the ability of α cells to adapt and respond properly to amino acid levels.

### Glutamine metabolism is required for maximal glucagon secretion

It has been well established that amino acids, including arginine and glutamine, stimulate glucagon secretion^21^. While arginine is a potent stimulator of glucagon secretion, glutamine alone does not stimulate glucagon secretion, but it is required for maximal glucagon secretion^33^. The mechanism by which glutamine affects glucagon secretion remains unknown, but glutamate has also been found to be co-secreted with glucagon and acts as a positive modulator for glucagon secretion through autocrine signaling^34^.

To understand how the loss of glutamine metabolism impacts α cell function, we exposed isolated islets from *Gls^fl/fl^*and α*Gls^KO^*mice to static conditions of varying concentrations of glucose and amino acids. We saw no significant differences in the glucagon content across all conditions and genotypes (Extended Figure 7A). The islets from the *Gls^fl/fl^* mice responded as expected with increased glucagon secretion in response to low glucose and arginine. The addition of glutamine potentiated the glucagon response (Extended Figure 7B). The loss of glutamine metabolism resulted in a blunted glucagon response to all secretagogues (Extended Figure 7B), indicating that glutamine metabolism plays a crucial role in α cell function.

To further characterize hormone secretion, we next assessed secretion dynamics using real-time measurements. To assess the dynamics of glucose response, we exposed isolated islets to high glucose and low glucose. This was then followed by exposure to arginine and a combination of arginine and glutamine all at euglycemia to assess the dynamics of amino acid response. Like the static secretion assay, we found no differences in glucagon content between the genotypes (Extended Figure 7C). We found that the loss of α cell-specific glutamine metabolism led to a blunted glucagon response to both glucose stimulation and amino acid stimulation (Figure 5A). Not only was there a ∼2.8-fold reduction in glucagon secretion in response to low glucose levels in α*Gls^KO^* islets (Figure 5B), but there was also a ∼2.3-fold reduction in basal glucagon secretion at high glucose levels (Figure 5C). Comparable results were observed in response to arginine (∼1.97-fold reduction) and arginine with glutamine (∼1.7-fold reduction) (Figure 5D-E). For both genotypes, we did not observe any sex-dependent differences in response to glucose or amino acids (Extended Figure 7D-O). Despite the significant effect on glucagon secretion, the loss of α cell-specific glutamine metabolism did not affect insulin content or secretion in response to glucose or amino acids under these conditions (Extended Figure 8A-F).

**Figure 5.**
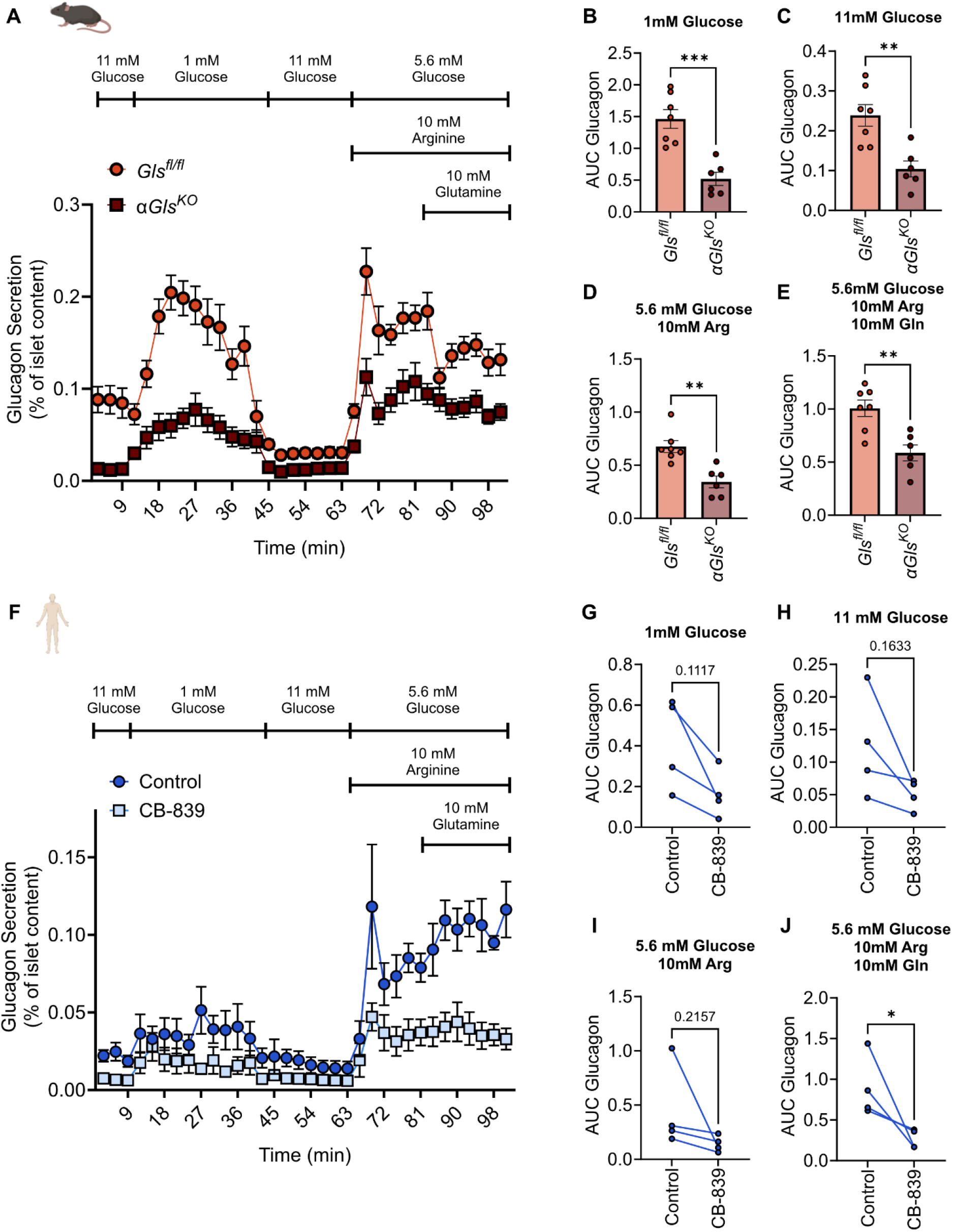
Glutamine metabolism is required for maximal glucagon secretion in mouse and human islets. (A) Islets from *Gls^fl/fl^* and α*Gls^KO^* were subjected to perifusion assays to assess real-time glucagon secretion in response to high (11 mM) glucose, low (1mM) glucose, and euglycemia (5.6 mM) in combination with arginine (10 mM) alone or arginine (10 mM) with glutamine (10 mM). (B) Area under the curve of the glucagon response to low (1 mM) glucose. (C) Area under the curve of the glucagon response to high (11 mM) glucose. (D) Area under the curve of the glucagon response to euglycemia (5.6 mM) in combination with arginine (10 mM) alone (E) Area under the curve of the glucagon response to euglycemia (5.6 mM) in combination with arginine (10 mM) and glutamine (10 mM). (F) Human cadaveric donor islets treated with either DMSO or CB-839 (100nM) were subjected to perifusion assays to assess real-time glucagon secretion in response to high (11 mM) glucose, low (1mM) glucose, and euglycemia (5.6 mM) in combination with arginine (10 mM) alone or arginine (10 mM) with glutamine (10 mM). (G) Area under the curve of the glucagon response to low (1 mM) glucose. (H) Area under the curve of the glucagon response to high (11 mM) glucose. (I) Area under the curve of the glucagon response to euglycemia (5.6 mM) in combination with arginine (10 mM) alone (J) Area under the curve of the glucagon response to euglycemia (5.6 mM) in combination with arginine (10 mM) and glutamine (10 mM).

### Glutamine metabolism supports human α cell response to amino acids

Given the implication that glutamine metabolism is altered in diabetes^35^, we assessed the effect of whole islet inhibition of glutaminase activity on human cadaveric donor islets. To do this, we treated human cadaveric donor islets with CB-839, or DMSO as a control for 24 hours. We then subjected these islets to a perifusion assay to assess real-time hormone secretion. We found that the inhibition of whole islet glutaminase activity did not impact glucagon content (Extended Figure 8G) but resulted in a decrease in glucagon secretion in response to amino acids, particularly arginine and glutamine in combination (Figure 5F). There was no significant difference between the control and CB-839 treated islets in response to glucose concentration or arginine (Figure 5G-I). Like the mouse islets, we see no significant differences in insulin secretion between the control and the treated islets (Extended Figure 8H-M).

## Discussion

Here, we identify glutaminase as a central regulator of α cell adaptation and function (Figure 6). Using an inducible α cell-specific *Gls* knockout model, we demonstrate that α cells constitute a major site of glutamine utilization within pancreatic islets and that glutaminase activity couples amino acid availability to both proliferative and secretory responses. These findings expand current models of islet nutrient sensing by identifying glutaminase as a key integrator of α cell growth, signaling, and hormone secretion.

**Figure 6.**
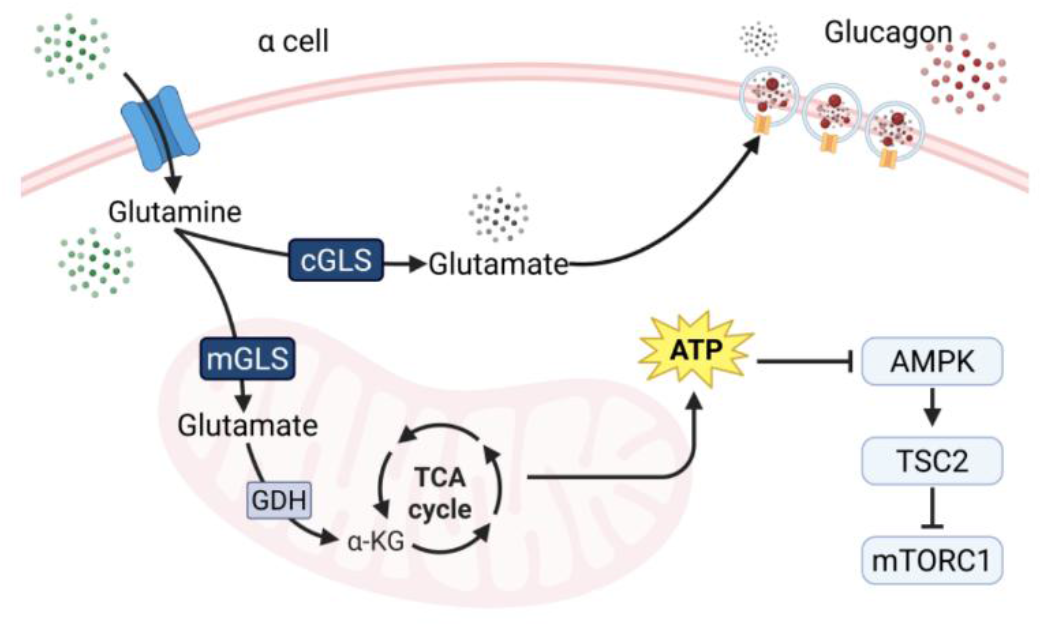
Schematic of the proposed mechanism by which glutaminase activity regulates α cell proliferation, mTORC1 activation and glucagon secretion.

Although glutaminase has long been recognized as the first step in glutamine catabolism, its role in α cell physiology remains poorly understood. We show that loss of glutaminase impairs amino acid-induced α cell proliferation, attenuates mTORC1 activation and mTORC2-AKT signaling, and diminishes glucagon secretion in response to both amino acid stimulation and hypoglycemia.

Our findings suggest that glutaminase serves a distinct role in α cells compared with β cells. Whereas glutamine metabolism primarily supports insulin secretion in β cells^36^, glutaminase coordinates α cell growth and glucagon secretion.

Glutamine metabolism in α cells has been linked to redox balance, AMPK signlaing^37^, and autocrine glutamatergic signaling that reinforces glucagon secretion^34^. Consistent with these observations, our findings support a model in which glutamine catabolism coordinates both metabolic and signaling pathways required for α cell function.

Importantly, we demonstrate that glutaminase-dependent regulation of α cell growth and glucagon secretion is conserved in human islets, underscoring the translational relevance of this pathway. Dysregulated amino acid metabolism, altered α cell function, and defective glucagon secretion are hallmarks of both type 1 and type 2 diabetes^38^. Consistent with the importance of adaptive α cell growth, germline loss-of-function mutations in the glucagon receptor that cause Mahvash disease are characterized by chronic hyperaminoacidemia, marked α cell hyperplasia, and pancreatic neuroendocrine tumor development, highlighting the potent proliferative effects of sustained amino acid signaling^39^. We further found that glutaminase expression in human α cells is positively correlated with HbA1C levels (R^2^ ∼0.66, P-value ∼8.5e-6) and T2D diabetic status (R^2^ ∼0.66, P-value ∼7.5e-6)^40^ with these findings mirrored in mice (T2D diabetic status: R^2^ ∼0.78, P-value ∼1.6e-7). Previous reports linking GLS genetic variation to islet gene expression^41,42^ and two SNPs that alter glutaminase expression in human islets (rs7563403 and rs6711082) are also associated with cardiovascular disease and circulating metabolite levels. Together, these findings raise the possibility that altered glutaminase activity contributes to defective α cell adaptation and impaired counterregulatory responses during metabolic disease.

Glutaminase-dependent nutrient sensing may enable α cells adapt to chronic alterations in nutrient availability, including the hyperaminoacidemia that accompanies insulin deficiency and glucagon resistance^4^. Defining how glutamine metabolism contributes to these adaptive responses may provide new insight into mechanisms underlying islet dysfunction in diabetes.

Interestingly, α cell-specific glutaminase deficiency also altered δ cell biology. Although the mechanism underlying this effect remains unknown, these findings raise the possibility that glutamine metabolism within α cells contributes to intra-islet communication. While functional coupling between β and δ cells through gap junctions has been well described^43^, communication between α and δ cells has received comparatively less attention. Emerging evidence suggests that a feedback loop between α and δ cells may play an important role in coordinating hormone secretion and cellular adaptation^44^. Our findings identify glutaminase-dependent α cell metabolism as a potential regulator of this signaling axis.

Despite establishing glutaminase as a critical regulator of α cell adaptation and function, there are several limitations that should be considered. Although glutaminase deficiency consistently impaired α cell proliferation, mTOR signaling, and glucagon secretion, the downstream mechanisms remain undefined and may involve glutamate production, mitochondrial metabolism, redox homeostasis, calcium signaling, or other nutrient-sensitive pathways. Furthermore, this study does not interrogate the previously defined signaling axis between mTOR and glucagon secretion^45^.

Although our findings were validated in human islets, the human studies relied on acute pharmacologic inhibition of glutaminase rather than genetic manipulation. In addition, all hormone secretion studies were performed in intact islets, limiting our ability to distinguish cell-autonomous effects from those mediated by α, β, and δ cell paracrine interactions. Since glucagon secretion is regulated by complex intra-islet signaling, and the organization and functional importance of these signaling networks differ between mouse and human islets^46^, our findings should be interpreted within this context. Therefore, because our secretion studies were not designed to specifically dissect α-to-β cell or α-to-δ cell interactions, we cannot exclude the possibility that loss of α cell-specific glutaminase alters endocrine function in part through changes in intra-islet paracrine signaling.

Future studies should define the molecular mechanisms by which glutaminase regulates α cell nutrient sensing and secretory competence. Defining how glutaminase regulates calcium signaling, cellular energetics, and redox homeostasis during amino acid stimulation and hypoglycemia will clarify how metabolic flux is translated into hormone secretion and adaptive cellular responses. Extending these studies to models of diet-induced obesity, insulin resistance, and diabetes will be essential to determine how α cell-specific glutaminase deficiency influences islet adaptation during metabolic disease and whether glutaminase-dependent nutrient sensing contributes to the compensatory α cell responses associated with type 2 diabetes. Finally, determining how glutaminase regulates intra-islet communication may reveal whether altered paracrine signaling, electrical coupling, or other forms of intercellular communication underlying the observed effects on δ cells.

Collectively, our findings establish glutaminase as a critical regulator of α cell nutrient sensing, adaptive growth, and endocrine function. More broadly, this work positions glutamine metabolism as a fundamental determinant of α cell adaptive capacity and expands current understanding of how pancreatic islets integrate nutrient availability with hormonal output. These results identify glutaminase as a previously unrecognized regulator of α cell biology and suggest that modulation of glutamine metabolism may represent a therapeutic strategy for disorders characterized by impaired islet adaptation and glucagon dysregulation.

## Acknowledgements

We thank Dr. Mark A Magnuson for valuable intellectual discussions and thoughtful insights that helped shape the direction and interpretation of this research.

This work used data acquired from the [Human Pancreas Analysis Program (HPAP)] (https://hpap.pmacs.upenn.edu/) (RRID:SCR_016202), supported by the Human Islet Research Network (HIRN; RRID:SCR_014393), with funding from UC4-DK112217, U01-DK123594, UC4-DK112232, and U01-DK123716. Additional islet samples and tissues were provided by the Integrated Islet Distribution Program (IIDP) (RRID:SCR_014387), funded by NIH Grant #2UC4DK098085.

Human pancreatic islets and/or other resources were provided by the NIDDK-funded Integrated Islet Distribution Program (IIDP) (RRID:SCR_014387) at City of Hope, NIH Grant # U24DK098085. Informed consent for deceased donors was obtained by the IIDP in accordance with the National Institute of Health.

This work includes data and/or analyses from HumanIslets.com funded by the Canadian Institutes of Health Research, JDRF Canada, and Diabetes Canada (5-SRA-2021-1149-S-B/TG 179092) with data from islets isolated by the Alberta Diabetes Institute Islet Core with the support of the Human Organ Procurement and Exchange program, Trillium Gift of Life Network, BC Transplant, Quebec Transplant, and other Canadian organ procurement organizations with written informed donor consent as approved by the Human Research Ethics Board at the University of Alberta (Pro00013094).

## Data Availability Statement

Human single cell RNA sequencing data is a publicly available dataset PanKbase^11^. Data can be found at: https://zenodo.org/records/15596314. Mouse genomic is a publicly available data set <u>GSE80673</u>^12^. Zebrafish genomic data is a publicly available data set available on ENA (http://www.ebi.ac.uk/ena) under accension number PRJEB10140^13^.

## Research Design and Methods

### Single Cell RNA Sequencing Analysis

Single cell RNA sequencing (scRNA-seq) data were obtained from the PanKBase dataset (version 3.3) and loaded into R (v4.2) using the Seurat package (v4.3) for downstream analysis. The dataset was stored as an .rds object and read into R using readRDS(). All analyses were performed using the default RNA assay, and cells were classified according to their assigned cell_classification identities as defined by PanKBase.

Genes involved in glutamine metabolism and transport were selected for targeted analysis. These included key metabolic enzymes (GLS, GLUL, GLUD1, OGDH, GOT1, GPX3, GCLC, GCLM, GLS2, ASNS, ASPG) and transporters (SLC38A5, SLC38A4).

For each glutamine transporter gene, expression data were extracted along with the corresponding cell type annotations using FetchData() and exported to CSV files for further inspection. Violin plots were generated using exported raw data and graphed in GraphPad Prism. Heat maps were generated based on mean gene expression from the exported data.

### Zebrafish Studies

The role of glutaminase in the regulation of α cell number in Zebrafish (Danio rerio) was studied using a previously described glucagon receptor double-knockout line expressing GFP under the control of the glucagon promoter to identify α cells (Tg(gcga:GFP);gcgra/b-/-)^47^. To knock down glsa and glsb, a pair of top-ranking sgRNAs (Extended Figure 9A), nominated by CHOPCHOP^48^, were synthesized for each gene as described^49^. The sgRNA pairs were mixed separately or in combination with recombinant Cas9 protein (PNA Bio Inc) at a 1.5:1 molar ratio with a final concentration of 1-5 µM of Cas9 and incubated at room temperature for 10 minutes. The resultant RNPs were injected into one cell stage embryos. The mutagenesis efficiency was assessed the next day using a heteroduplex formation assay^49^. Batches with >85% mutagenesis were used for total GFP+ α cell counting in the islets at five days post-fertilization (dpf) under a fluorescent microscope, as described previously.

### Mouse Studies

All mouse studies were performed at Vanderbilt University Medical Center and approved by the Institutional Animal Care and Use Committee. Mice were housed on a 12:12 hour light:dark cycle with *ad libtum* access to standard rodent chow, unless indicated for fasting purposes, and water. GLS1^fl/fl^ mice were generated from Gls^tm1a(KOMP)Mbp^ embryonic stem cells (Project ID: CSD29307), which were used for blastocyst microinjection to produce chimeric animals (Duke University Transgenic and Knockout Shared Resource). Following FLP-mediated excision of the selection cassette, GLS1^fl/fl^ mice were crossed with Gcg-Cre^ERT2^ transgenic mice (MMRRC Strain #042277-JAX), in which a tamoxifen-inducible Cre recombinase is expressed under the control of the glucagon (*Gcg*) promoter, to generate α cell-specific GLS1 knockout animals. Mice were injected with 100uL of a 20mg/mL tamoxifen solution in corn oil once a day for 3 days. Following a 2-week washout period, mice were treated weekly with 10mg/kg of a humanized monoclonal antibody targeting the glucagon receptor (mAb) intraperitoneally once a week for 2 weeks to interrupt glucagon receptor signaling. Blood glucose was monitored every 3 days for 14 days post-treatment period via tail vein nick and measured using a Contour Next-EZ hand-held glucometer. Both male and female mice were combined in these studies as there were no differences were observed between sexes.

The presence of Cre was verified through immunostaining (methods described below) of Cre (Cell Signaling, 15036S, 1:500) and glucagon to mark α cells. The knockout of *Gls* was verified through immunostaining of GLS1 (Protein Tech, 12855-1-AP, 1:250), KGA (Protein Tech, 20170-1-AP, 1:250), and GAC (Protein Tech, 19958-1-AP, 1:250). Quantification was conducted using CellCounter via Image J. Percent of Cre+, GLS1+, KGA+, or GAC+ α cells were calculated from at least 500 α cells per animal by dividing the co-positive glucagon+ cells by total glucagon+ cells.

### Metabolic Tolerance Tests

For serum hormone levels, mice were fasted for ∼6 hours and then intraperitoneally injected with 2mg arginine/kg body weight. Blood was obtained via the retroorbital sinus vein from isoflurane anesthetized mice using a heparinized glass capillary tube (Fisherbrand, 22-260-950) and immediately transferred to a 1.5mL Eppendorf tubes containing a protease inhibitor cocktail (Invitrogen D1306). Samples were left on ice for 45 minutes and then were centrifuged 2000g for 10 minutes at 4⁰C. Serum was collected from the blood samples and stored at -80⁰C. Insulin (Catalog no. 10-1247-01, Mercodia Inc) and glucagon (Catalog no. 10-1281-01, Mercodia Inc) levels were measured by ELISA.

For mixed meal tolerance tests, mice were fasted for ∼6 hours and administered a bolus of original vanilla ensure by oral gavage (Abbott SKU#: 57243) (10μL/80% of b.w. in grams). Blood glucose was measured from tail-bleeds at the following points: 0, 15, 30, 45, 60, 90, 120 minutes.

For glucose tolerance tests, mice were fasted for ∼6 hours and treated with a 2mg/kg glucose bolus via intraperitoneal injection. Blood glucose was measured from tail-bleeds at the following points: 0, 15, 30, 45, 60, 90, 120 minutes.

For insulin tolerance tests, mice were fasted for ∼6 hours and treated with 0.075 U/ml insulin (recombinant human insulin, no. I9278; Sigma-Aldrich) in filter-sterilized 1× PBS at 0.1 ml/10 g body weight via intraperitoneal injection. Blood glucose was measured from tail-bleeds at the following points: 0, 15, 30, 45, 60, 90, 120 minutes.

### Islet Isolation from Mice

Pancreatic islets were isolated by dissecting the splenic portion of the pancreas and injecting collagenase P (Millipore Sigma Roche, 11249002001) (0.618mg/mL) into the bile duct. The digestion was then aided by using a wrist-action shaker for 10 minutes. The islets were then collected via histopaque (Sigma-Aldrich, 10771) (1.077g/mL) gradient separation. Islets were resuspended in RPMI media supplemented with 10%FBS, 0.01% Penicillin Streptomycin, and 5.6mM glucose. Islets were handpicked to near 100% purification through consecutive rounds in islet RPMI media and were cultured overnight in islet RPMI media or used immediately for experiments.

### Human Pancreas and Islet Studies

For immunofluorescent staining, deidentified human pancreas samples were also obtained through IIDP. Deidentified human donor information is provided in Extended Figure 9B-C. Islets were cultured in CMRL containing HI-FBS, 1M HEPES, GlutaMAX, Pen/Strep, and glucose for 24 hours and then treated with CB-839 (100nM) or DMSO for an additional 24 hours.

### Time-Course Glutaminase Activity *Ex Vivo*

Islets were isolated from *Gls^fl/fl^* and α*Gls^KO^* mice. The islets were lysed in Tris-HCL lysis buffer (10mM Tris-HCL, 150mM NaCl, 1.0% Triton-X, 1mM EDTA, 1X Halt™ Protease Inhibitor Cocktail, pH 8.6) on ice. Upon extraction of cell debris, the supernatant was then subjected to pre-warmed 5mM glutamine in Tris-HCL lysis buffer. The lysate was incubated at 37°C and heat quenched at 95°C at 0, 15, 30, 45, and 60 minutes. Any remaining debris was removed via centrifugation. Endogenous glutamate production was measured using the Glutamine/Glutamate-Glo™ Assay (Promega, J8022). Glutaminase activity was calculated by the slope of the glutamate production over time.

### Tissue Collection from Mice

Upon the completion of the treatment period, the pancreata and blood serum samples were collected. Blood serum was left on ice for 45 minutes and spun down at 2000g for 10 min at 4°C. Serum was collected from the blood samples and stored at -80⁰C. Histology samples were fixed in 4% PFA and washed with 10X PBS. After subjecting the tissue to a 30% sucrose bath, they were embedded in OCT (Fisher Scientific, 23-730-571) and stored at -80°C until use.

### Immunofluorescence

Islets were isolated from 14- to 16-week-old mice as described above. These islets were then cultured in media containing high amino acids, low amino acids, high amino acids without glutamine, or high amino acids with CB-839 for 5 days. After culture, islets were washed in 2mM EDTA and dispersed with 0.025% Trypsin at 37°C for 10 minutes with mixing. Dispersed islet cells were recovered by centrifugation in RPMI media containing 5.6mM glucose, 10% FBS, and 1% Penicillin/Streptomycin. The resulting cell pellet was resuspended in medium and centrifuged onto a glass slide using a Cytospin^TM^ 4 (Thermo Scientific, Waltham, MA) centrifuge. The slides were then dried at room temperature overnight and stored at -80°C until use. The slides were then thawed, fixed in 4% paraformaldehyde, and stained for glucagon (LSBio, LS-C202759, 1:1000) and Ki67 (Abcam, ab15580, 1:500). The percentage of proliferating α cells was calculated based on the number of Ki67^+^/glucagon^+^ cells divided by the total number of glucagon^+^ cells using Imaris image analysis software (Oxford Instruments). This methodology was repeated on human islets. Pancreas from 14- to 16-week-old mice were prepared for histology as described above. Whole mouse pancreas was sectioned on a cryostat at a thickness of 8μm per section. The full depth of the pancreas was sectioned in 5 μm increments by collecting 10 sections and then removing 7 sections. This process was repeated until 7 depths of 150 μm was collected from each pancreas. For islet cell mass analysis, one section from each of the seven depths was stained for C-peptide (Invitrogen, PA-85595, 1:500) to mark β cells, glucagon to mark α cells and somatostatin (Invitrogen, MA5-16987, 1:250) to mark δ cells. Whole pancreatic sections were imaged on Leica DMi8 inverted microscope (Leica Microsystems) under 10X magnification and image analysis was conducted via ImageJ (FIJI). Total pancreatic islet cell masses were calculated as previously described^50^. Islet cell areas from each of seven depths were normalized to total pancreas section area. Areas from each of the seven sections were summed and the sum multiplied by total pancreas mass to achieve an estimate of the percent of total mass. Islet cell areas were then compared from a midplane section of the pancreas to investigate islet size.

For α cell proliferation analysis, pancreas form mice treated for two weeks with GCGR mAb or IgG were sectioned and immunostained for the proliferation marker, Ki67, and glucagon to mark α cells. Islets were imaged on Leica DMi8 inverted microscope (Leica Microsystems) and co-positive cells were counted using CellCounter via Image J. Percent α cell proliferation was calculated from at least 500 α cells per animal by dividing the Ki67^+^/glucagon^+^ cells by total glucagon^+^ cells. The same analysis was repeated for phospho-S6 (240/244) (Cell Signaling, 5364S, 1:500), phospho-S6 (235/236) (Cell Signaling, 2211S, 1:500), cleaved caspase 3 (Asp175) (Biotechne, MAB835, 1:500), Phospho-AMPKα (Thr172) (Cell Signaling, 2531S, 1:500), Phospho-Tuberin/TSC2 (Ser1387) (Cell Signaling, 5584S, 1:500), and SLC38A5 (Sigma, SAB2105557, 1:500).

For α cell number per islet, the number of α cells were counted per islet for 30 islets in each pancreas. For α cell area, the total area of the glucagon stain of an image was calculated using ImageJ. The total area was then normalized to the number of α cells in the image. This was repeated until at least 500 α cells were quantified.

### Hormone Secretion Assays

For *in vitro* static glucagon secretion assays, islets were isolated from mice and equilibrated in RPMI containing 0.5 mg/mL BSA supplemented with 10% FBS and 5.5 mM glucose for 1 h at 37°C, 5% CO2. ∼20 islets/well were picked into 500mL RPMI at glucose concentrations specified in the Figure in 24-well plate(s) and glucagon secretion was measured over 1 h at 37°C, 5% CO2. Supernatants were then collected, supplemented with 1X Halt™ Protease Inhibitor Cocktail (1:100), and stored at −20°C until analysis. Glucagon secretion was measured using Mouse Glucagon ELISA kit (Mercodia, 10-1281-01).

Dynamic glucagon and insulin secretion were measured from islet chambers loaded with ∼100 islets/chamber (BioRep Technologies) and perifused at a flowrate of 100uL/min in response to glucose and amino acid stimulations indicated in the figure legends. The effluent was collected at 3 min-intervals. Glucagon concentration in each perifusion fraction and total glucagon content was measured by a glucagon ELISA assay (Mercodia, 10-1281-01). Insulin concentration in each perifusion fraction and total insulin content was measured by an insulin ELISA assay (Mercodia, 10-1247-01).

### Statistical Analysis

All data are shown with error bars indicating standard error of the mean. Each point represents an individual animal. Data within individual experiments were compared with ordinary two-way ANOVA using Tukey correction for multiple comparisons, unless otherwise designated in the figure legend.

---

**Extended Figure 1.**
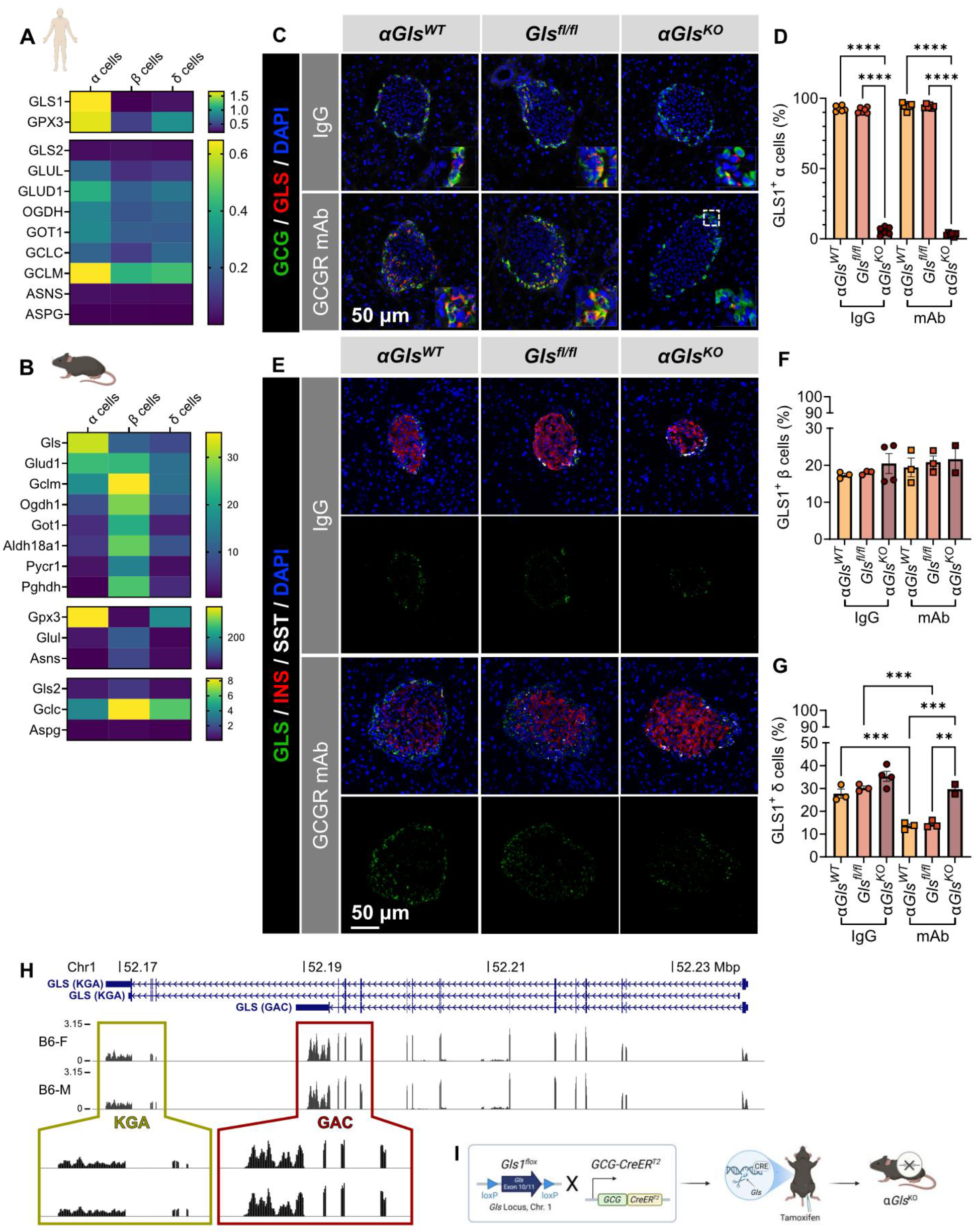
(A) Heat map representing mean gene expression of glutamine metabolism genes in human α, β, and δ cells. (B) Heat map representing mean gene expression of glutamine metabolism genes in mouse α, β, and δ cells. (C) Representative immunostaining for GCG and GLS1 from αGls^WT^, Glsfl^/fl^, and αGls^KO^ mice (scale bar = 50 μm). (D) Quantification of GLS1 in α cells as determined by percent GLS1+/GCG+ cells per total GCG+ cells in α*Gls^WT^, Glsfl^/fl^*, and α*Gls^KO^* mouse islets after two weeks of treatment with GCGR mAb or control IgG (n=6 each). (E) Representative immunostaining for GLS1, C-peptide, and somatostatin from αGls^WT^, Glsfl^/fl^, and αGls^KO^ mice treated with 2 weeks of IgG or mAb (scale bar = 50 μm). (F) Quantification of GLS1 in β cells as determined by percent GLS1+/INS+ cells per total INS+ cells in α*Gls^WT^, Glsfl^/fl^*, and α*Gls^KO^* mouse islets after two weeks of treatment with GCGR mAb or control IgG (n=3 each). (G) Quantification of GLS1 in δ cells as determined by percent GLS1+/SST+ cells per total SST+ cells in α*Gls^WT^, Glsfl^/fl^*, and α*Gls^KO^* mouse islets after two weeks of treatment with GCGR mAb or control IgG (n=3 each). (H) Differential exon usage analysis of KGA (ENSMUST00000114513, ENSMUST00000114512) and GAC (ENSMUST00000114510) in C57BL/6 mice (I) Schematic representation of the mouse breeding scheme to generate α*Gls^KO^* mice

**Extended Figure 2.**
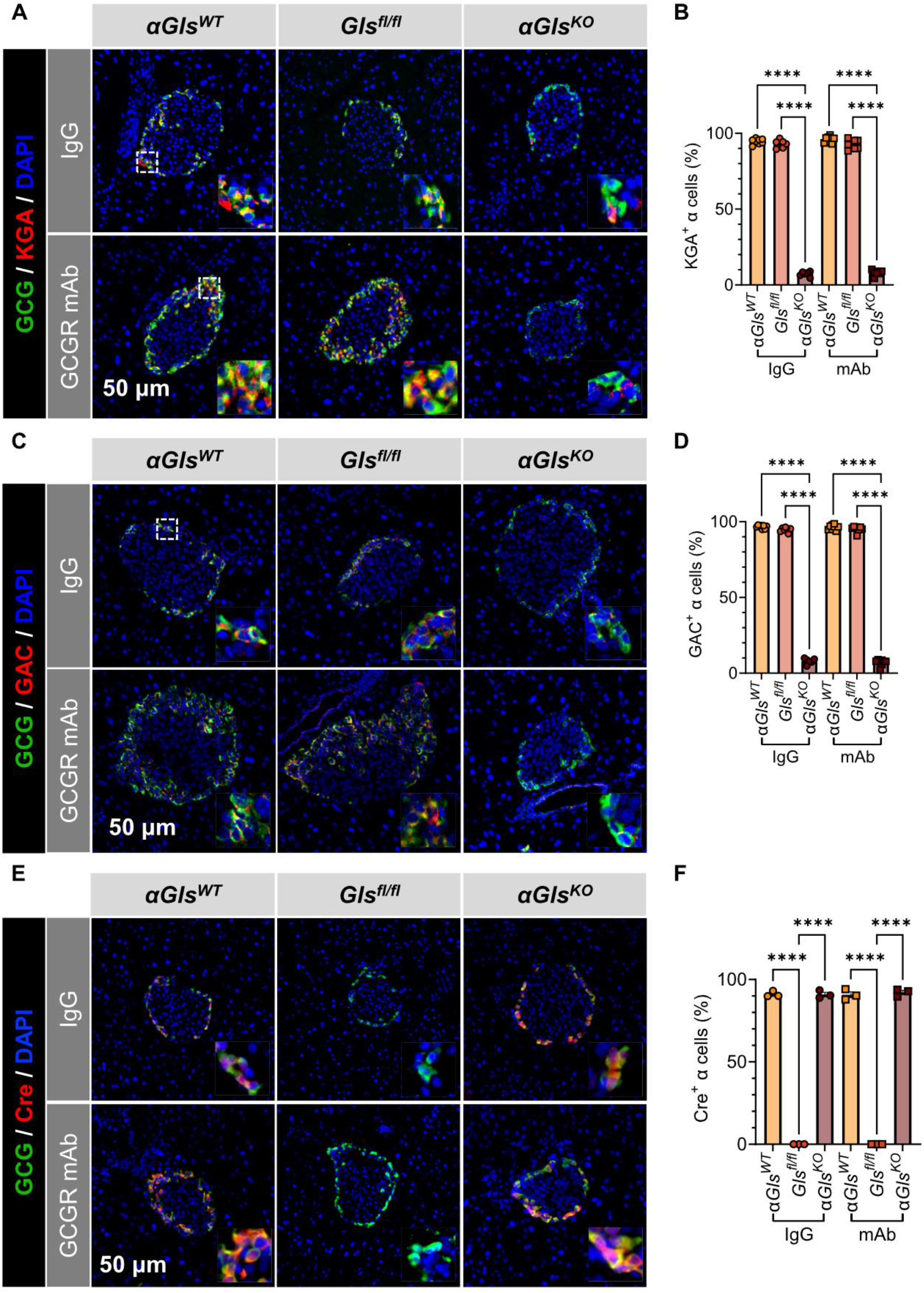
(A) Representative immunostaining for glucagon and KGA from α*Gls^WT^, Glsfl^/fl^*, and α*Gls^KO^* mice (scale bar = 50μm, inset width = 22 μm). (B) Quantification of KGA expression in α cells as determined by KGA^+^/GCG^+^ cells per total GCG^+^ cells in α*Gls^WT^, Glsfl^/fl^*, and α*Gls^KO^* mouse islets after treatment with GCGR mAb or control IgG (n=6-8). (C) Representative immunostaining for glucagon and GAC from α*Gls^WT^, Glsfl^/fl^*, and α*Gls^KO^* mice (scale bar = 50μm, inset width = 22 μm). (D) Quantification of GAC expression in α cells as determined by GAC^+^/GCG^+^ cells per total GCG^+^ cells in α*Gls^WT^, Glsfl^/fl^*, and α*Gls^KO^* mouse islets after treatment with GCGR mAb or control IgG (n=6-8). (E) Representative immunostaining for glucagon and Cre from αGls^WT^, Glsfl^/fl^, and αGls^KO^ mice (scale bar = 50 μm, inset width = 22 μm). (F) Quantification of Cre expression in α cells as determined by percent Cre+/GCG+ cells per total GCG+ cells in αGls^WT^, Glsfl^/fl^, and αGls^KO^ mouse islets after two weeks of treatment with GCGR mAb or control IgG (n=3 each).

**Extended Figure 3.**
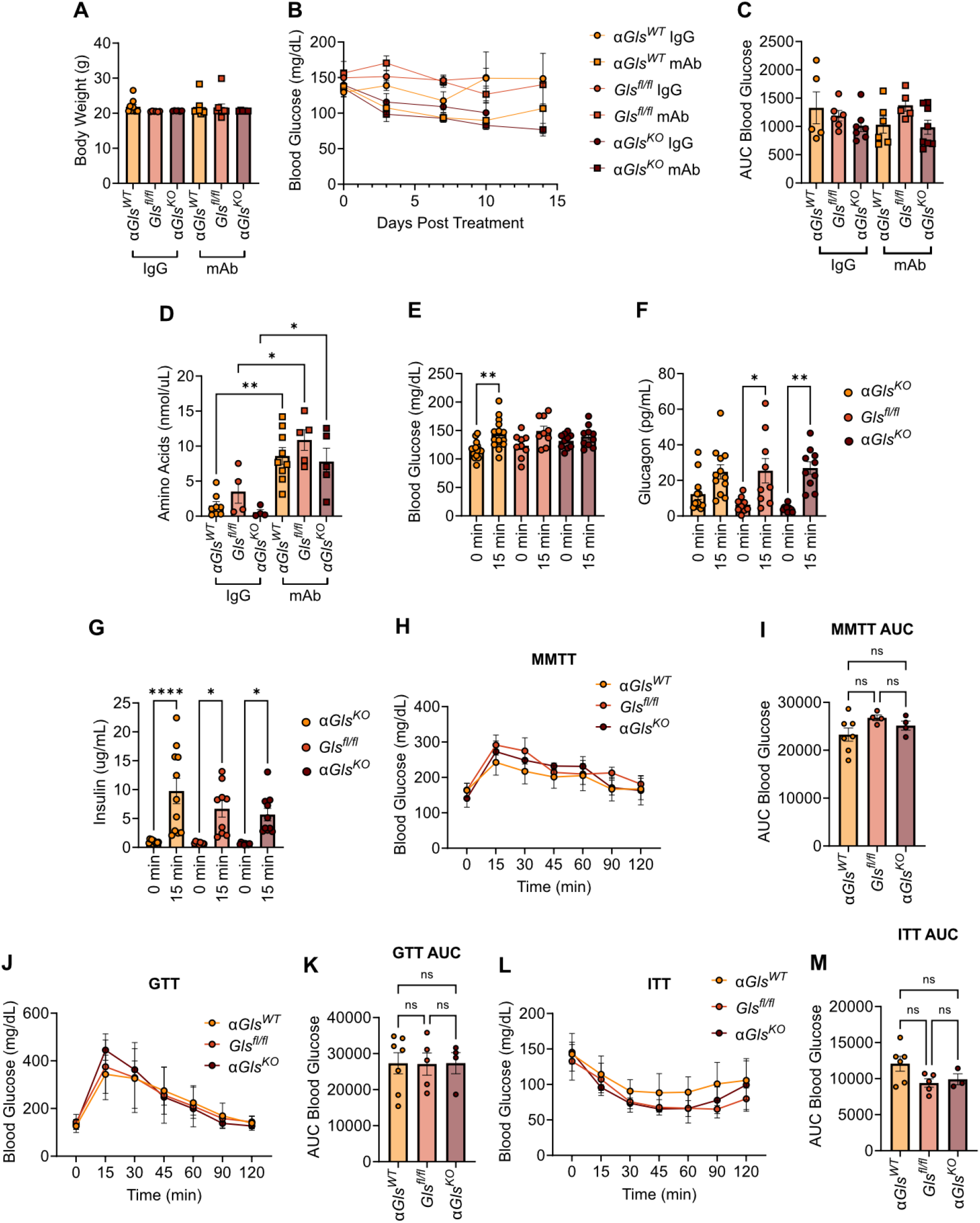
(A) Body weight of α*Gls^WT^, Glsfl^/fl^*, and α*Gls^KO^* mice. (B) Blood glucose was measured every 3 days for 2 weeks post antibody treatment. (C) Calculated area under the curve of blood glucose over 2 weeks post antibody treatment (n=5-7). (D) Circulating amino acid concentrations 2 weeks post antibody treatment (n=4-8). (E) Arginine stimulated blood glucose measured at 0 minutes and 15 minutes. (F) Arginine stimulated serum glucagon at 0 minutes and 15 minutes. (G) Arginine stimulated serum insulin at 0 minutes and 15 minutes (H) Blood glucose over 2 hours in response to a mixed meal tolerance test. (I) Area under the curve of blood glucose in mixed meal tolerance test. (J) Blood glucose over 2 hours in response to a glucose tolerance test. (K) Area under the curve of blood glucose in glucose tolerance test. (L) Blood glucose measured over 2 hours in response to an insulin tolerance test. (M) Area under the curve of blood glucose in insulin tolerance test.

**Extended Figure 4.**
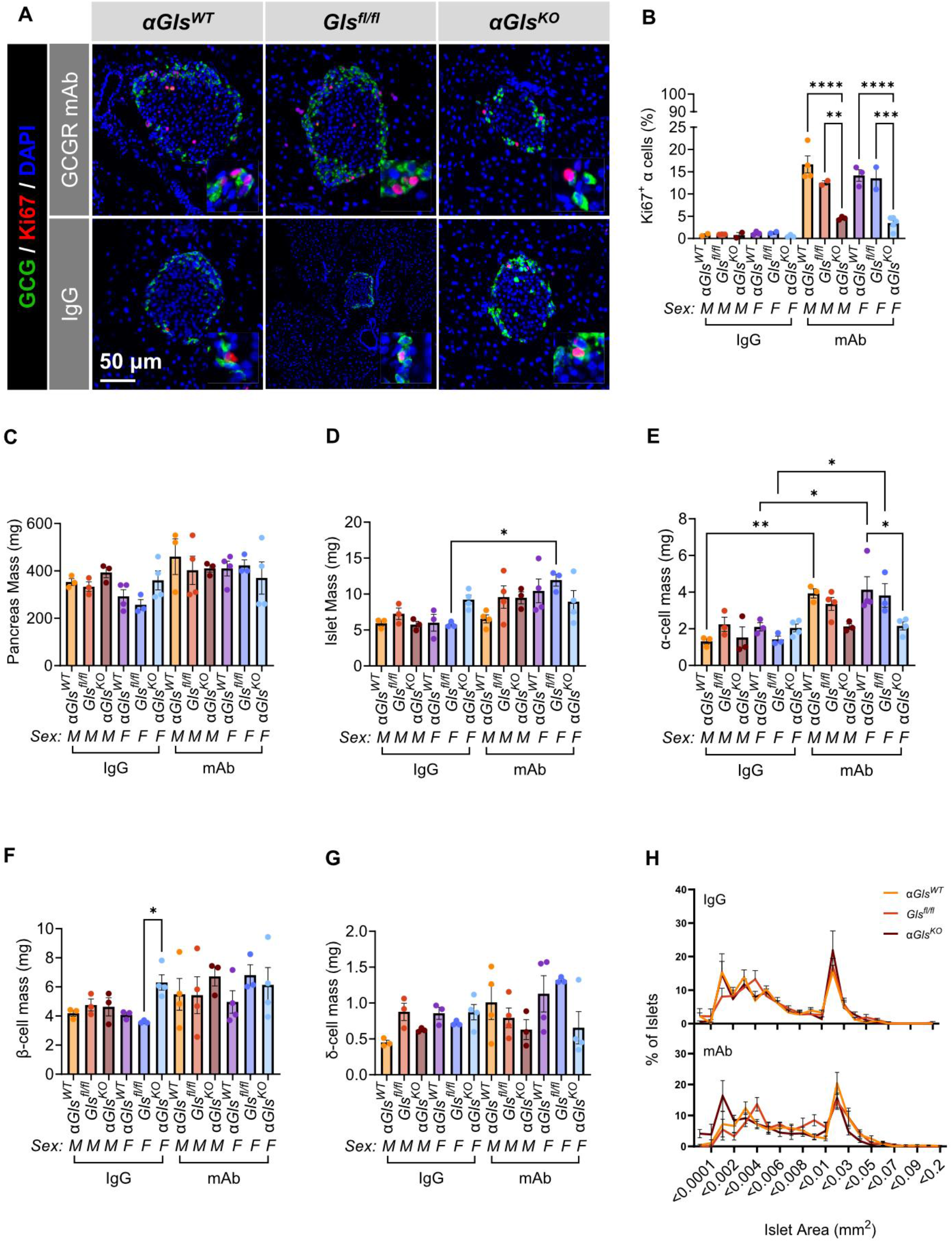
(A) Representative immunostaining for glucagon and Ki67 from α*Gls^WT^, Glsfl^/fl^*, and α*Gls^KO^* mice (scale bar = 50 μm, inset width = 22 μm). (B) Analysis of α cell proliferation as indicated by Ki67 by sex. (C) Percentage of total islets by islet area. (D-H) Analysis of pancreas mass (D), islet mass (E), α cell mass (F), β cell mass (G), and δ cell mass (H) by sex.

**Extended Figure 5.**
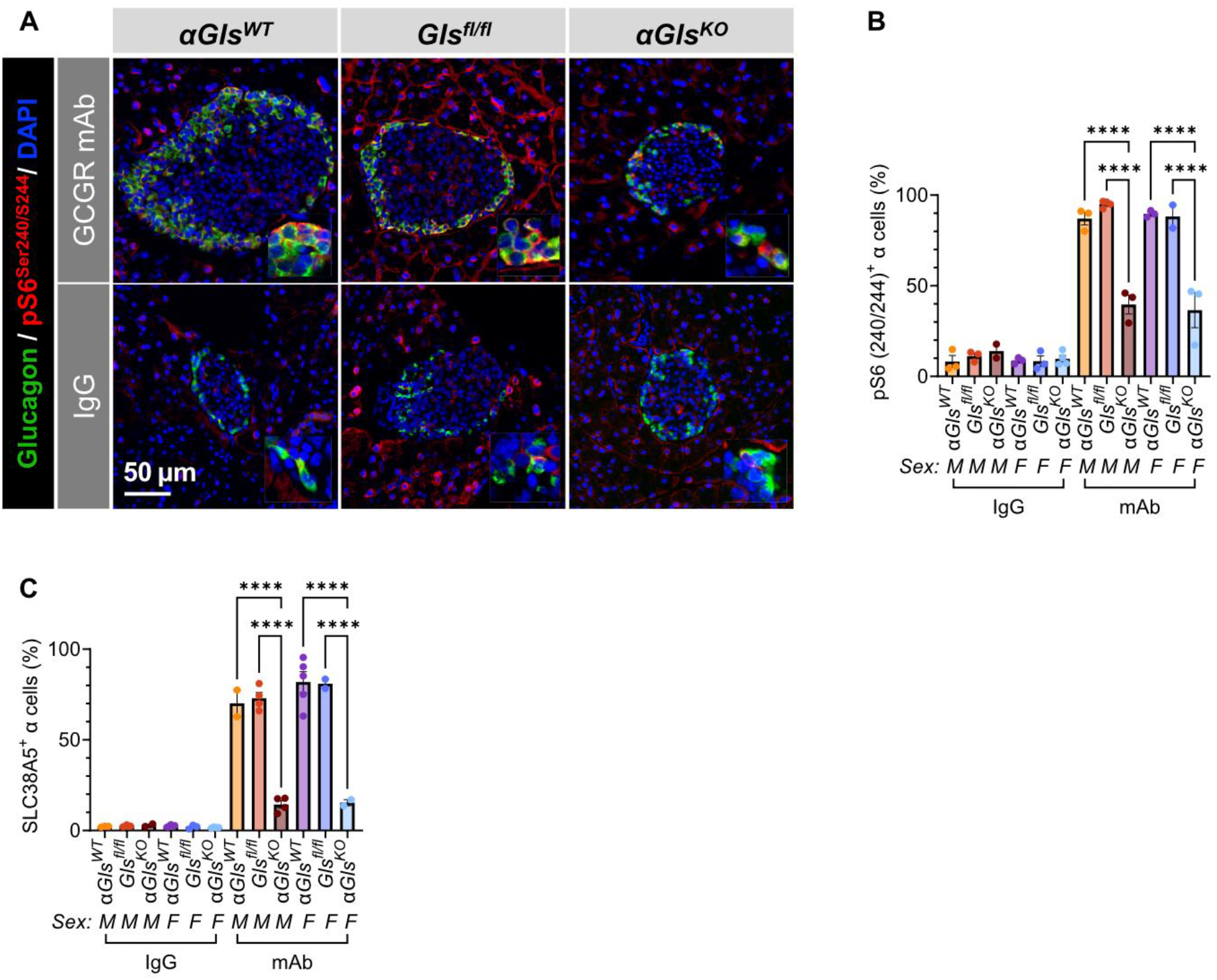
(A) Representative immunostaining for glucagon and pS6(240/244) from α*Gls^WT^, Glsfl^/fl^*, and α*Gls^KO^* mice (scale bar = 50 μm, inset width = 22 μm). (B) Analysis of mTORC1 activation in α cells as indicated by pS6(240/244) by sex. (C) Analysis of SLC38A5 expression in α cells by sex.

**Extended Figure 6.**
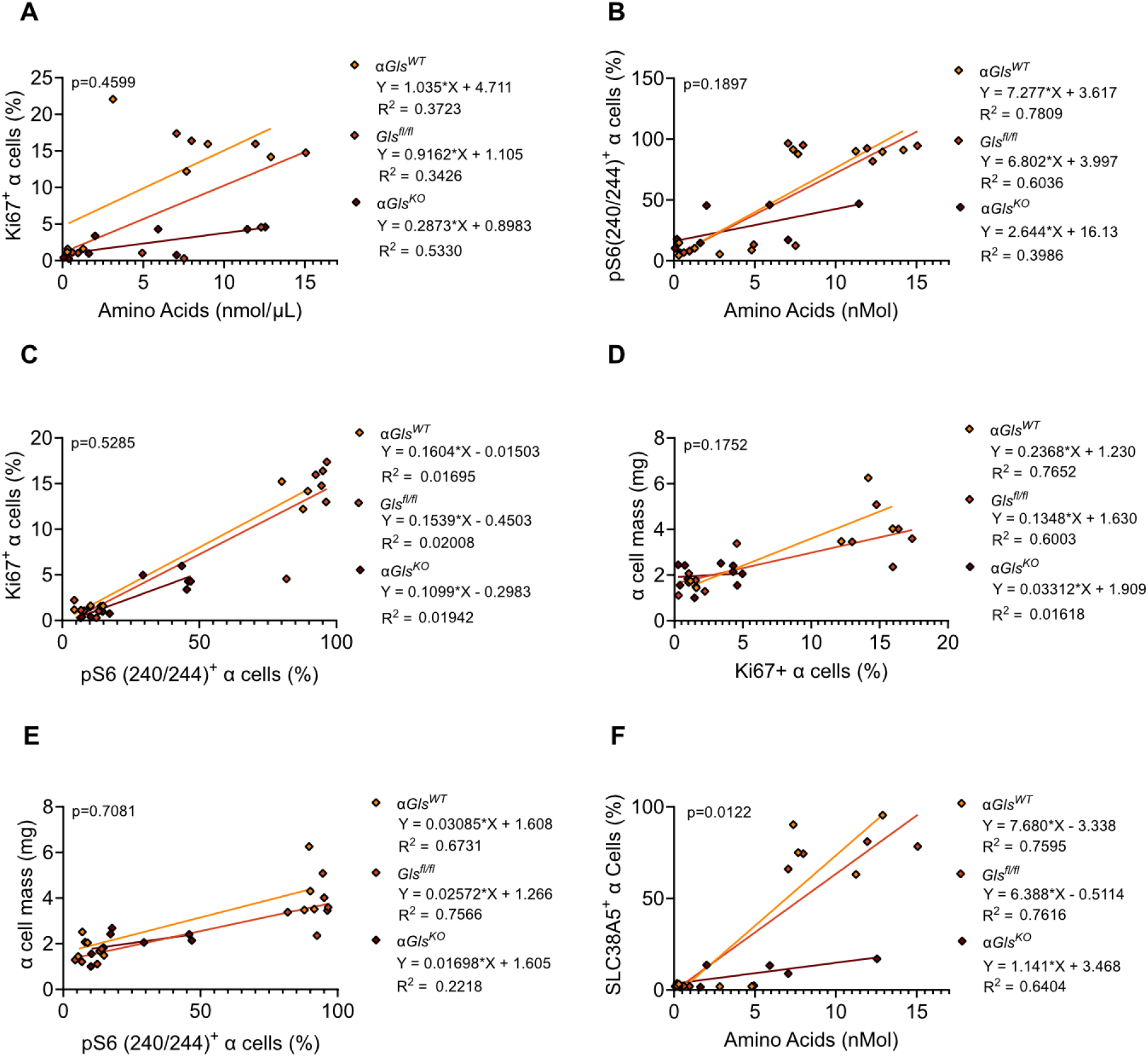
(A) Simple linear regression of circulating amino acids and proliferating α cells (Ki67) from α*Gls^WT^, Glsfl^/fl^*, and α*Gls^KO^* mice treated with either IgG or mAb. (B) Simple linear regression of circulating amino acids and α cells with mTORC1 activation (pS6) from α*Gls^WT^, Glsfl^/fl^*, and α*Gls^KO^* mice treated with either IgG or mAb. (C) Simple linear regression of proliferating α cells and α cells with mTORC1 activation (pS6) from *Gls^WT^, Glsfl^/fl^*, and α*Gls^KO^* mice treated with either IgG or mAb. (D) Simple linear regression of proliferating α cells and α cell mass from α*Gls^WT^, Glsfl^/fl^*, and α*Gls^KO^* mice treated with either IgG or mAb. (E) Simple linear regression of α cells with mTORC1 activation (pS6) and α cell mass from α*Gls^WT^, Glsfl^/fl^*, and α*Gls^KO^* mice treated with either IgG or mAb. (F) Simple linear regression of circulating amino acids and SLC38A5 expression in α cells from α*Gls^WT^, Glsfl^/fl^*, and α*Gls^KO^* mice treated with either IgG or mAb.

**Extended Figure 7.**
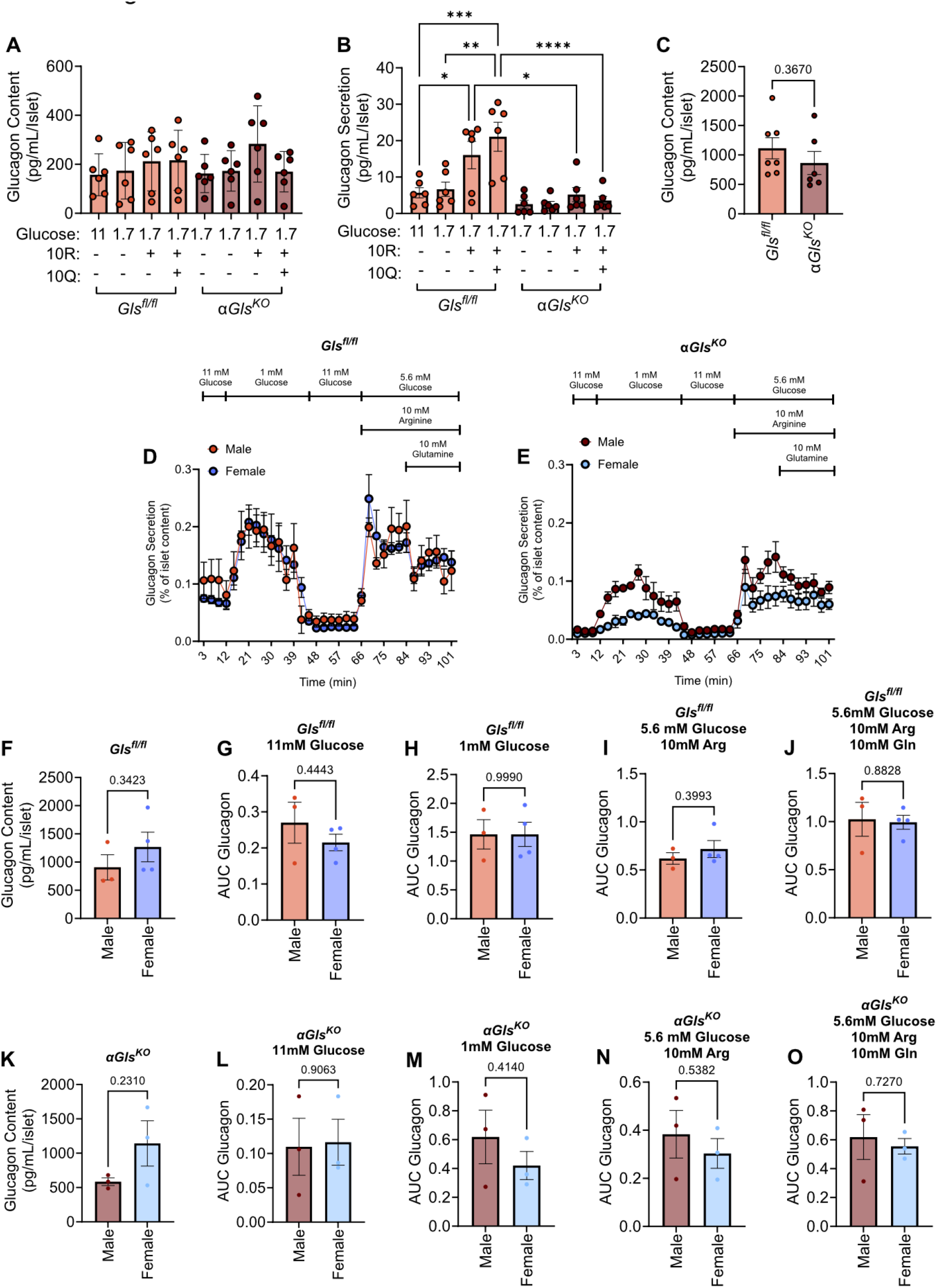
(A) Glucagon content of *Gls^fl/fl^* and α*Gls^KO^* mouse islets when exposed to high (11 mM) glucose or low (1 mM) glucose in combination with arginine (10 mM) alone or arginine (10mM) with glutamine (10 mM). (B) Static secretion assay normalized to the number of islets of *Glsfl^/fl^* and α*Gls^KO^* mouse islets when exposed to high (11 mM) glucose or low (1 mM) glucose in combination with arginine (10mM) alone or arginine (10 mM) with glutamine (10 mM). (C) Glucagon content of *Gls^fl/fl^* and α*Gls^KO^* mouse islets subjected to perifusion assay. (D) Islets from *Gls^fl/fl^* mice were subjected to perifusion assays to assess real-time glucagon secretion in response to high (11 mM) glucose, low (1mM) glucose, and euglycemia (5.6 mM) in combination with arginine (10 mM) alone or arginine (10 mM) with glutamine (10 mM) by sex. (E) Islets from α*Gls^KO^* mice were subjected to perifusion assays to assess real-time glucagon secretion in response to high (11 mM) glucose, low (1mM) glucose, and euglycemia (5.6 mM) in combination with arginine (10 mM) alone or arginine (10 mM) with glutamine (10 mM) by sex. (F) Glucagon content from *Gls^fl/fl^* islets by sex. (G-J) Area under the curve of the glucagon response from *Gls^fl/fl^* islets to low (1 mM) glucose (G), high (11 mM) glucose (H), and euglycemia (5.6 mM) in combination with arginine (10 mM) alone (I) or arginine (10 mM) and glutamine (10 mM) (J) by sex. (K) Glucagon content from α*Gls^KO^* islets by sex. (L-O) Area under the curve of the glucagon response from α*Gls^KO^* islets to low (1 mM) glucose (L), high (11 mM) glucose (M), and euglycemia (5.6 mM) in combination with arginine (10 mM) alone (N) or arginine (10 mM) and glutamine (10 mM) (O) by sex.

**Extended Figure 8.**
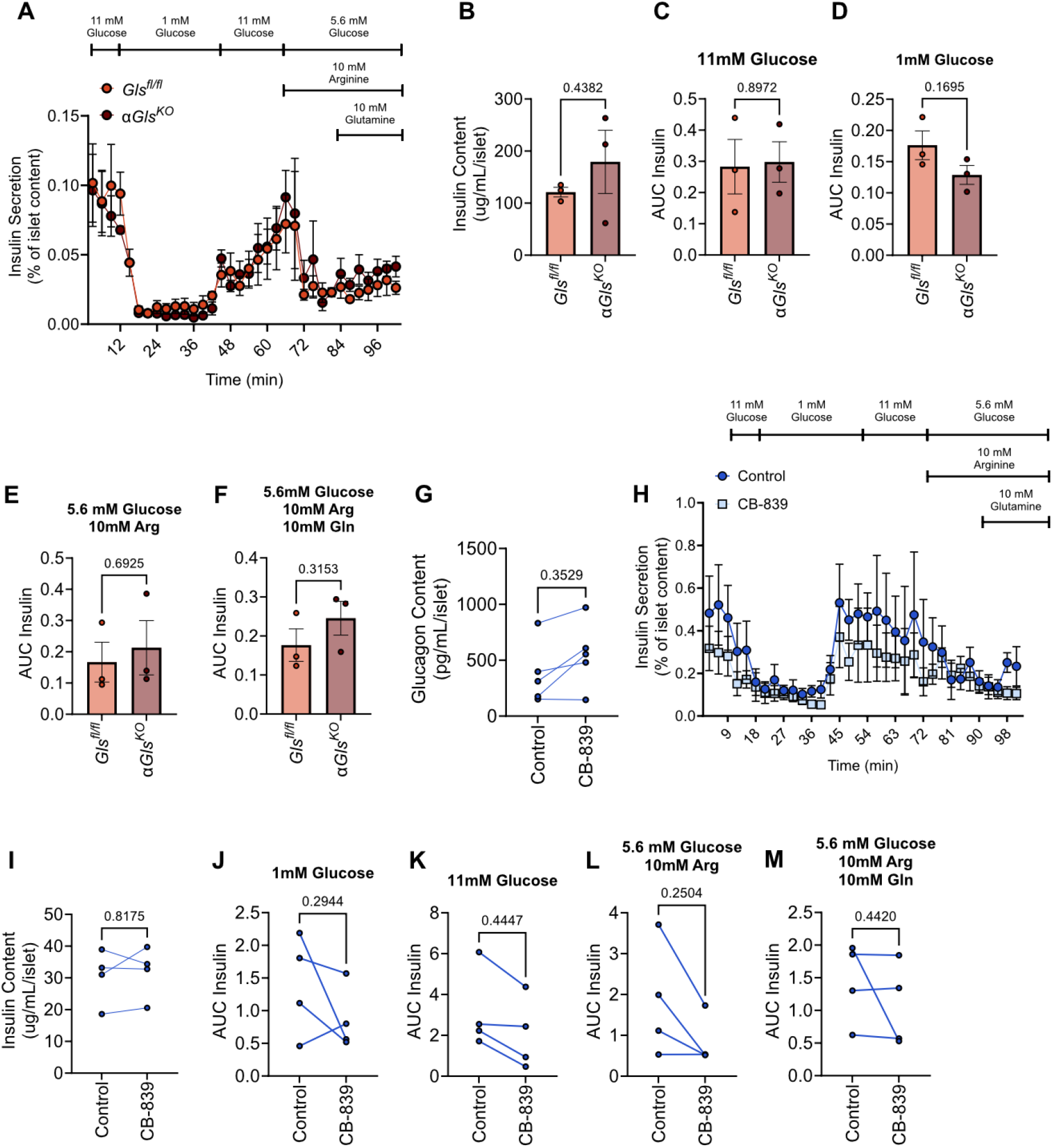
(A) Islets from *Gls^fl/fl^* and α*Gls^KO^* mice were subjected to perifusion assays to assess real-time insulin secretion in response to high (11 mM) glucose, low (1mM) glucose, and euglycemia (5.6 mM) in combination with arginine (10 mM) alone or arginine (10 mM) with glutamine (10 mM) by sex. (B) Insulin content from *Gls^fl/fl^* and α*Gls^KO^* islets. (C-F) Area under the curve of the insulin response from *Gls^fl/fl^* and α*Gls^KO^* islets to low (1 mM) glucose (C), high (11 mM) glucose (D), and euglycemia (5.6 mM) in combination with arginine (10 mM) alone (E) or arginine (10 mM) and glutamine (10 mM) (F). (G) Glucagon content of human cadaveric donor islets treated with DMSO or CB-839 subjected to perifusion assay. (H) Human cadaveric donor islets treated with DMSO or CB-839 were subjected to perifusion assays to assess real-time insulin secretion in response to high (11 mM) glucose, low (1mM) glucose, and euglycemia (5.6 mM) in combination with arginine (10 mM) alone or arginine (10 mM) with glutamine (10 mM) by sex. (I) Insulin content from human cadaveric donor islets treated with DMSO or CB-839. (J-M) Area under the curve of the insulin response from human cadaveric donor islets treated with DMSO or CB-839 to low (1 mM) glucose (J), high (11 mM) glucose (K), and euglycemia (5.6 mM) in combination with arginine (10 mM) alone (L) or arginine (10 mM) and glutamine (10 mM) (M).

**Extended Figure 9.**
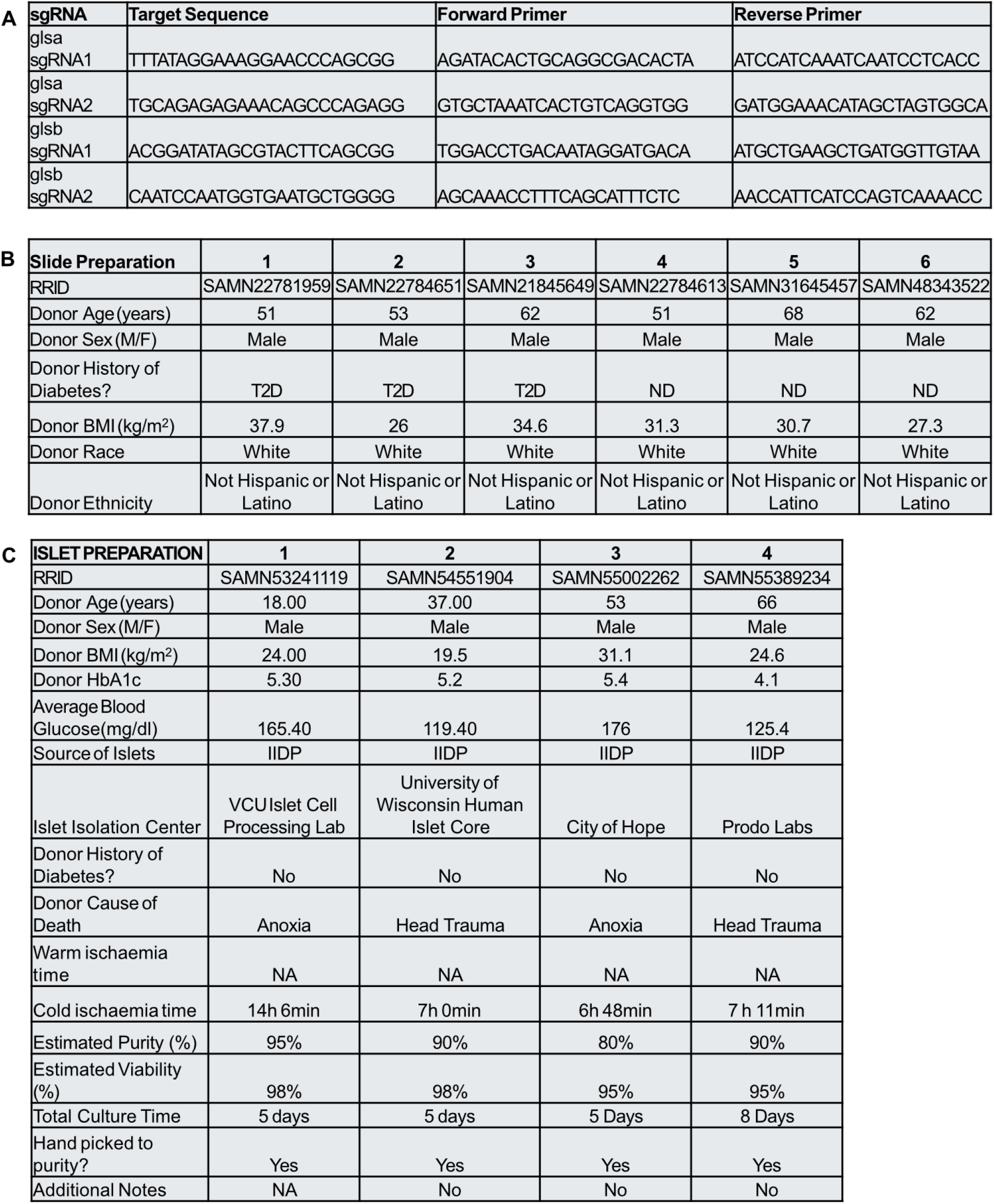
(A) Donor characteristics for human pancreas slides. (B) Donor characteristics for cadaveric human islets. (C) Primer sequences for glsa and glsb knockdown in zebrafish via sgRNA.

